# ZODIAC: database-independent molecular formula annotation using Gibbs sampling reveals unknown small molecules

**DOI:** 10.1101/842740

**Authors:** Marcus Ludwig, Louis-Félix Nothias, Kai Dührkop, Irina Koester, Markus Fleischauer, Martin A. Hoffmann, Daniel Petras, Fernando Vargas, Mustafa Morsy, Lihini Aluwihare, Pieter C. Dorrestein, Sebastian Böcker

**Affiliations:** Chair for Bioinformatics, Friedrich-Schiller-University, Jena, Germany; Collaborative Mass Spectrometry Innovation Center, Skaggs School of Pharmacy and Pharmaceutical Sciences, University of California San Diego, La Jolla, USA; Skaggs School of Pharmacy and Pharmaceutical Sciences, University of California San Diego, La Jolla, USA; Scripps Institution of Oceanography, University of California San Diego, La Jolla, USA; International Max Planck Research School “Exploration of Ecological Interactions with Molecular and Chemical Techniques”, Max Planck Institute for Chemical Ecology, Jena, Germany; Division of Biological Science, University of California San Diego, La Jolla, USA; Department of Biological and Environmental Sciences, University of West Alabama, Livingston, USA

## Abstract

The confident high-throughput identification of small molecules remains one of the most challenging tasks in mass spectrometry-based metabolomics. SIRIUS has become a powerful tool for the interpretation of tandem mass spectra, and shows outstanding performance for identifying the molecular formula of a query compound, being the first step of structure identification. Nevertheless, the identification of both molecular formulas for large compounds above 500 Daltons and novel *molecular formulas* remains highly challenging. Here, we present ZODIAC, a network-based algorithm for the *de novo* estimation of molecular formulas. ZODIAC reranks SIRIUS’ molecular formula candidates, combining fragmentation tree computation with Bayesian statistics using Gibbs sampling. Through careful algorithm engineering, ZODIAC’s Gibbs sampling is very swift in practice. ZODIAC decreases incorrect annotations 16.2-fold on a challenging plant extract dataset with most compounds above 700 Dalton; we then show improvements on four additional, diverse datasets. Our analysis led to the discovery of compounds with novel molecular formulas such as C_24_H_47_BrNO_8_P which, as of today, is not present in any publicly available molecular structure databases.

## 2 Introduction

Metabolomics characterizes metabolites with high-throughput techniques. In the past decade, liquid chromatography coupled with tandem mass spectrometry (LC-MS/MS) became widely adopted by the metabolomics community as a powerful and sensitive analytical platform, but yet only a fraction of the detected compounds can be effectively annotated, even partially. In untargeted LC-MS/MS experiments, thousands of metabolites can be detected and fragmented from a single biological sample. The measured fragmentation spectra (MS/MS spectra) are then used to examine the metabolite’s structure. But confident high-throughput annotation of those MS/MS spectra remains highly challenging; this prevents the building of comprehensive knowledge in fields where metabolomics has become central, such as biomedical research, natural products drug discovery^1^, environmental science, and food science. Tandem mass spectrometry data is often searched against spectral libraries^2^; unfortunately, spectral libraries are of limited size, in particular when it comes to biomolecules^3^. Although public spectral libraries are being expanded, only a fraction of acquired MS/MS spectra can be annotated by spectral library searches^1,4-6^.

Computational methods have been developed that do not search in spectral libraries but rather in molecular structure databases^7^. Among these methods, CSI:FingerID^8^ has repeatedly shown the best performance^9-12^. One reason for CSI:FingerID’s improved performance is the integration of SIRIUS^10^, deducing the molecular formula of each query as the first step of its analysis. Other tools filter candidates using the query precursor mass, reducing molecular formula annotation to a “byproduct”. This worsens identification rates^11^ and can result in severe “hidden prior” problems^13,14^. Identifying the molecular formula is also the very first step in structural elucidation using Nuclear Magnetic Resonance (NMR) or X-ray crystallography, guiding data interpretation based on atoms and unsaturation degree. The confident annotation of molecular formulas from mass spectrometry data is far from trivial, especially if executed *de novo* (without a structure database): Here, the number of candidate molecular formula grows rapidly with the compound size and elements beyond CHNOPS. To counter this growth, one can use heuristic constraints^15^ or use only molecular formulas from some structure database^16,17^. Restricting the search space will improve the performance of a method in evaluation, but will prevent the discovery of novel molecular formulas in application.

Arguably the best-performing computational method for molecular formula annotation is SIRIUS 4^10^, which combines isotope pattern matching^15,18-23^ and MS/MS fragmentation tree computation^22,24-26^. SIRIUS reaches best-of-class performance without filtering or meta-scores^24^. But even SIRIUS has problems annotating molecular formula for compounds above 500 Da: Böcker & Dührkop^24^ found that the percentage of correctly identified molecular formulas dropped substantially for larger masses.

An alternative approach to annotate molecular formulas for a complete LC-MS run uses Gibbs sampling and Bayesian statistics, utilizing co-occurrence of molecular formulas differing by a predefined set of biotransformations^27-30^. Implicitly, these approaches try to identify molecular structures (or their isomers) from a restricted structure database, and cannot annotate novel molecular formulas. Network visualization approaches which connect compounds by hypothetical biotransformations and common chemical functional groups have been demonstrated to ease manual molecular formula annotation^31^. Independently, network-based methods were developed for structural elucidation and dereplication^1,32,33^. All of these approaches are based on the fact that compounds in an LC-MS run usually co-occur in a network of derivatives.

## 3 Results and Discussion

We present ZODIAC (ZODIAC: Organic compound Determination by Integral Assignment of elemental Compositions) for confident, database-independent molecular formula annotation in LC-MS/MS data. ZODIAC takes advantage of the fact that an organism produces related metabolites that are derived from multiple, but limited, biosynthetic pathways. ZODIAC builds upon SIRIUS and uses, say, the top 50 molecular formula annotations from SIRIUS as candidates for one compound. ZODIAC then reranks molecular formula candidates using Bayesian statistics. Prior probabilities are derived from fragmentation tree similarity, which supports reciprocal plausibility within an LC-MS/MS dataset. On the theoretical side, we establish that finding an optimal solution to the resulting computational problem is non-deterministic polynomial time (NP)-hard; to this end, we resort to Gibbs sampling. Using extensive algorithm engineering, Gibbs sampling running times were reduced to a practical level. To boost robustness, ZODIAC can integrate spectral library search hits. We show that ZODIAC improves molecular formula annotation on a diverse set of biological samples. Furthermore, ZODIAC scores allow us to rank molecular formula annotations by confidence. ZODIAC is not limited to molecular formulas from some structure database, allowing us to discover novel molecular formulas not present in any structural databases.

ZODIAC was evaluated on five diverse datasets representing samples from plants, human plasma, marine microalgae and mice fecal sample, see Supplementary Table 1, 5 and 6 and Supplementary Fig. 6. Input mzML/mzXML files were processed with OpenMS^34^ and low-quality MS/MS spectra were discarded, see Supplementary Fig. 7, Supplementary Table 2 and 7 and Section Materials & Methods. We evaluated SIRIUS and ZODIAC against a ground truth which was established by spectral library search and manual validation.

For all five datasets, we observe that ZODIAC outperforms SIRIUS, often substantially decreasing molecular formula annotation error rates (Fig. 1, left). We first consider the dendroides dataset, for which improvements are most distinctive: This dataset contains many larger compounds, and 75 % of the ground truth compounds have an *m/z* of 605 or higher (Supplementary Fig. 6). Hence, this dataset is particularly challenging for molecular formula assignment. Out of the 201 ground truth compounds, the preprocessing assigned an incorrect adduct to three; for these, the correct molecular formula is not contained in the candidate list considered by ZODIAC. For one compound, the corrected molecular formula was not ranked into the top 50. For the remaining 197 compounds, SIRIUS correctly annotated 50.76% (100), compared to 96.95% (191) for ZODIAC without anchors. This represents an 16.17-fold decrease in error rate. Error rates improves for compounds over the whole mass range, see Fig. 1 (right).

**Fig. 1:**
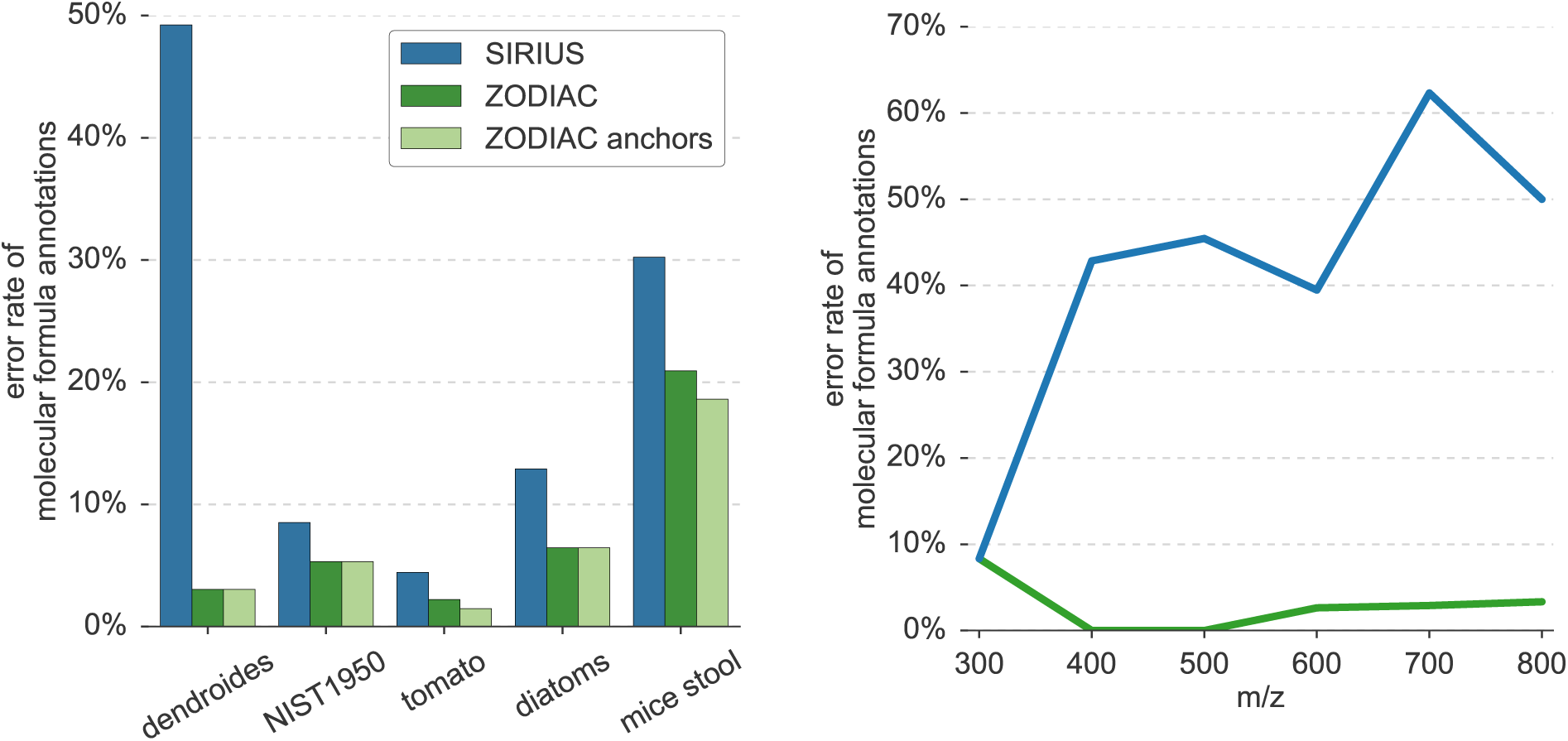
Molecular formula annotation error rates. Left: Error rate on five datasets. The rate of incorrect molecular formula annotations is displayed for SIRIUS and ZODIAC, with and without anchors. For number of compounds and other statistics, see Supplementary Table 1. ZODIAC reduces error rates on all datasets, but largest improvements are observed for the dendroides dataset which contains challenging, high molecular mass compounds. Right: Error rates vs. mass on dendroides dataset. Error rates for SIRIUS and ZODIAC without anchors are binned by compound *m/z*; bins of width 100 are centered at 300, 400, …, 800 *m/z*.

On the NIST1950 and tomato datasets, SIRIUS already showed excellent performance, with more than 90% correctly annotated molecular formulas. ZODIAC further decreased error rates, from 8.51% to 5.32% for NIST1950 and 4.44% to 2.22% for tomato. The diatoms dataset is rather complex, and many compounds may contain halogens and silicon. Here, SIRIUS reaches an error rate of 12.90%, which is reduced two-fold to 6.45% by ZODIAC.

Mice stool MS/MS spectra were measured with a broader isolation window, and the dataset contains numerous chimeric and low quality spectra, most of which were discarded before running ZODIAC. Consequently, ZODIAC has a much smaller network of interdependent compounds than for the other datasets. But even spectra that were not discarded often have substantially worse quality than spectra from other datasets: For example, these spectra often contain isotope peaks of fragments or are undetected chimeric spectra. Our evaluation shows that even for this extremely challenging dataset, ZODIAC improves annotation results, decreasing the error rate from 30.23% for SIRIUS to 20.93% for ZODIAC.

Some compounds in an LC-MS/MS run can result in high-scoring hits when searching in a MS/MS spectral library. ZODIAC’s stochastic model allows us to integrate these hits as *anchors*, assuming that we can trust assigned molecular formulas to a high degree. We performed a 10-fold cross-validation to assess the improvement using anchors. We ensured structure-disjoint evaluation on the library hits, as multiple “compounds” in the dataset may correspond to the same structure; see ref. ^8^ on the importance of structure-disjoint evaluation. ZODIAC with anchors improves the error rate on the tomato dataset to 1.48% and on mice stool to 18.60%, but does not improve results for the other datasets.

For four datasets, ground truth molecular formulas were established by library searching only. We tested if there is a distinct difference between the cosine score of ZODIAC’s correct and incorrect molecular formula assignments, but did not find such a difference (Supplementary Fig. 8).

We find that differentiating between adducts [M+H]^+^ and [M+Na]^+^ is sometimes challenging for SIRIUS and ZODIAC. This is observable for the dendroides dataset, where all six incorrect ZODIAC annotations show an incorrect adduct annotation, mistaking [M + H]^+^ for [M + Na]^+^ or vice versa. In all six cases, the molecular formula of the best ZODIAC hit and the ground truth differ by exactly two carbon minus two hydrogen atoms (21.984349 Da), with mass difference highly similar to that between [M + H]^+^ and [M + Na]^+^ (21.981944 Da). Sodium-ionized compounds can produce protonated fragments, making the interpretation of these spectra challenging. We reran ZODIAC on the dendroides dataset, assuming we knew the correct adduct for each reference compound. For all 201 compounds, the correct hit is contained in the SIRIUS top 50 candidate list. SIRIUS correctly annotated 67.17% (135) and ZODIAC 99.50% (200) of the compounds, corresponding to a 66-fold decrease of the error rate. The other four datasets contain fewer sodium adducts.

ZODIAC implicitly tries to estimate the probability that an annotated molecular formula is correct; as expected from the statistical theory, we find these estimates to be imprecise, see Fig. 2. But the ZODIAC score can be used to differentiate between true and incorrect annotations: For each dataset, we sort molecular formula annotations by the ZODIAC score, and calculate the rate of correct annotations for any subset of top-scoring annotations. We find that high-scoring ZODIAC annotations are more likely to be correct, see again Fig. 2. For this evaluation, we also considered previously discarded compounds for which SIRIUS did not rank the correct molecular formula in the top 50; for these compounds, ZODIAC cannot find the correct molecular formula but at best, the incorrect molecular formula should receive low ZODIAC scores. Selecting a ZODIAC score threshold of 0.9 results in more than 96.5% correct annotations while keeping 52.05% to 88.24% of the compounds of each dataset (Fig. 2). In comparison, spectral library search allowed us to annotate between 3.78% and 16.55% of a dataset, see Supplementary Table 1.

**Fig. 2:**
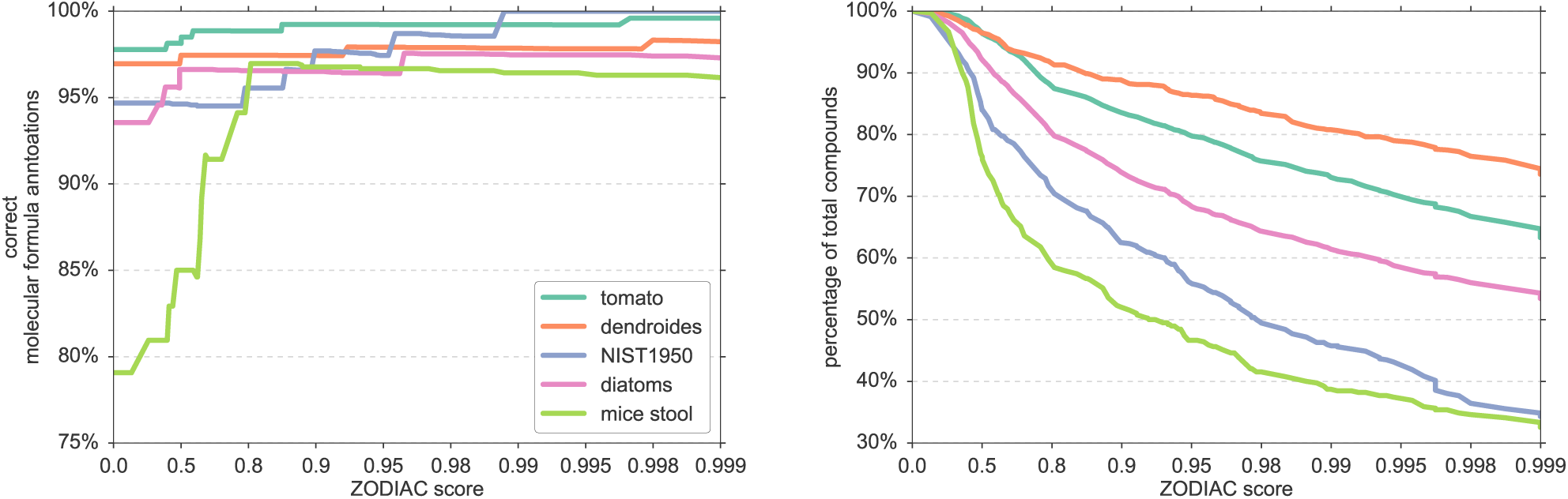
Percentage of correct annotations and number of compounds in relation to ZODIAC score. Left: Percentage of correct molecular formula annotations for different ZODIAC score thresholds for five datasets. We sort compounds by ZODIAC score and calculate the rate of correct annotations for all compounds above the given thresholds. Right: Percentage of total compounds with a ZODIAC score above different thresholds on five datasets. Here, we consider *all* compounds, with and without established ground truth. Note that scores on the x-axis are not equidistant.

### Novel molecular formulas

We now concentrate on *novel* molecular formulas, in the sense that these molecular formulas are not contained in the largest public molecular structure databases PubChem^17^ and ChemSpider^35^. As detailed in Section Materials & Methods, we cannot rule out that the molecular formula corresponds to, say, the compound minus a water loss instead of the full compound. Clearly, the structure of any compound with a novel molecular formula is also absent from the structure databases.

We use strict filters for both the ZODIAC score, the quality of the underlying MS/MS data, and the support by other molecular formulas in the dataset. We report molecular formula annotations from all five datasets with a) minimum ZODIAC score of 0.999, b) at least 95% of the MS/MS spectrum intensity being explained by SIRIUS, and c) at least one molecular formula of the compound is connected to 20 or more compounds in the ZODIAC similarity network. The third criterion discards compounds where ZODIAC’s results are basically identical to SIRIUS’s. This results in three novel molecular formulas in the tomato dataset and 59 in the diatoms dataset, see Supplementary Table 3 and 4. Filtering less restrictively (ZODIAC score at least 0.99, at least 90% of the MS/MS spectrum intensity being explained by SIRIUS), we annotate 31 in tomato, 103 novel molecular formulas in the diatoms dataset, one in NIST1950 and one in mice stool.

It is understood that some of these annotations may be wrong; unfortunately, a complete evaluation would require a full structural elucidation, which is experimentally infeasible. But our results clearly show that ZODIAC allows the user to select a few, potentially highly interesting compounds from a set of hundreds or thousands with low effort. Furthermore, using an example, we show in the next section that one top-scoring annotation from Supplementary Table 3 is presumably correct.

### Detailed evaluation of a novel bromine-containing molecular formula

We now concentrate on one particular compound in the diatoms dataset (*m/z* 588.230, retention time 503.97 sec): This ZODIAC-annotated compound is protonated and has molecular formula C_24_H_47_BrNO_8_P, which is indeed absent from the structure databases. The occurrence of bromine agrees with our expectation that marine organisms can be prolific sources of organohalogens^36^. The ZODIAC score of this annotation is 1.0, the maximum value. We found multiple lines of evidence that this molecular formula annotation is correct, both in the measured isotope pattern and the three MS/MS spectra measured for *m/z* 588.230 (presumably the monoisotopic peak of the isotope pattern), 590.228 (presumably the M+2 peak) and 592.325 (presumably the M+4 peak), see Fig. 3. To annotate fragments with molecular formulas, we used SIRIUS to compute a fragmentation tree for the MS/MS spectrum of the monoisotopic peak at *m/z* 588.230 (Fig. 3e).

**Fig. 3:**
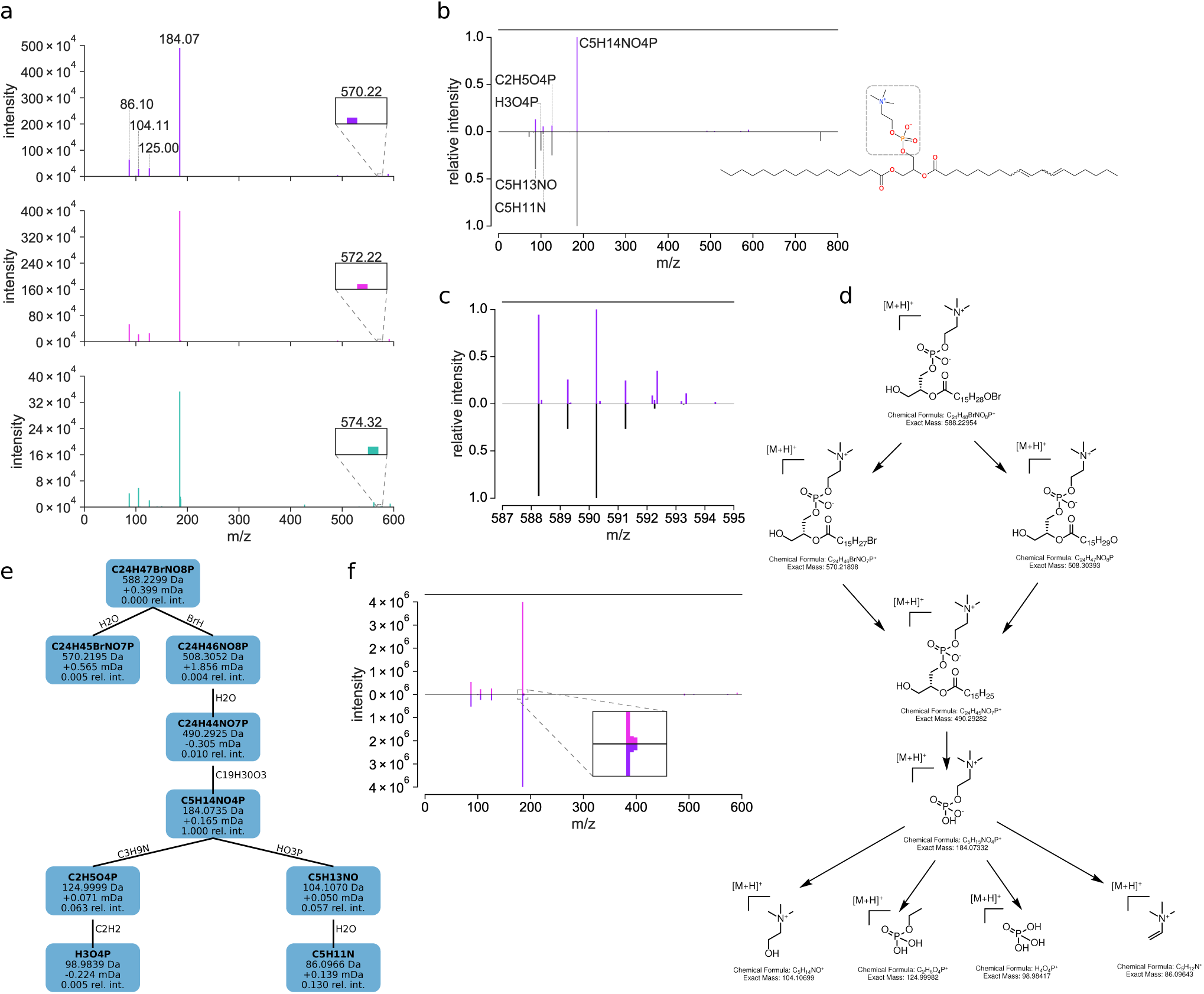
Annotation of a novel bromine-containing compound in the diatoms dataset. (a) MS/MS spectra for *m/z* 588.230, 590.228 and 592.325, corresponding to the monoisotopic, the M+2 and M+4 peak. The “moving” peak at *m/z* 570, 572 and 574 corresponds to the same molecular formula but different isotopes. This annotated fragment molecular formula is based on the fragmentation tree in (e) and is the only one containing bromine in these MS/MS spectra. (b) Partial match to 1-Palmitoyl-2-linoleoyl-sn-glycero-3-phosphocholine in the NIST library. The mirror plot compares the MS/MS spectrum of the monoisotopic peak at *m/z* 588.230 (top) to the NIST library spectrum (bottom). Displayed molecular formulas were annotated using the fragmentation tree of the query compound (e), and are identically annotated in the NIST reference spectrum. The substructure of the NIST reference compound which corresponds to the annotated peaks is highlighted. (c) Mirror plot of measured against simulated isotope pattern. The top part displays *m/z* 587 to 595 of the MS1 spectrum with retention time 503.97 sec, measured prior to the MS/MS spectrum for precursor *m/z* 588.230. The bottom part is the simulated isotope pattern for [C H BrNO P+ H]^+^. (d) Putative structure and fragmentation pathway of the novel compound. (e) Fragmentation tree computed by SIRIUS. Nodes correspond to fragments, edges to neutral losses. Nodes are annotated with the (neutralized) molecular formula, peak *m/z*, mass deviation in mDa and relative intensity. (f) Mirror plot of measured (top) against simulated (bottom) MS/MS spectrum for precursor M+2.

1. We compared MS/MS spectra for *m/z* 588.230, 590.228 and 592.325 (Fig. 3a) and found them to be highly similar, confirming that these peaks are indeed isotope peaks of one compound. One peak “moves” between MS/MS spectra (nominal *m/z* 570, 572 and 574): This peak corresponds to the fragment with molecular formula C_24_H_45_BrNO_7_P, which is the only annotated fragment containing bromine.
2. The measured isotope pattern agrees well with the theoretical isotope pattern of [C_24_H_47_BrNO_8_P + H]^+^ (Fig. 3c). The M+2 peak of the measured isotope pattern has a relative intensity of 106.0% of the monoisotopic peak, which is characteristic for the presence of a bromine atom.
3. The MS/MS spectrum for *m/z* 588.230 contains a precursor loss of 79.925 Da, and the only possible molecular formula explanation of this loss is BrH, considering a mass error of 100 ppm and elements CHNOPSFIClBrNaKSi.
4. We can simulate an MS/MS spectrum of the M+2 *peak that includes isotope patterns of fragments:* We use peak intensities from the MS/MS spectrum of the monoisotopic peak, and simulate isotope patterns of fragments as described in ref. ^37^. This allows us to verify whether the isotope patterns of fragments agree with our theoretical expectations. Indeed, simulated and measured MS/MS spectra of the M+2 peak show very high similarity, see Fig. 3f. The MS/MS spectrum of the M+4 peak must be treated with caution, as the precursor’s intensity is much lower and a second compound of higher intensity is present within the isolation window, see Supplementary Fig. 10. With regards to the moving peak, we can observe matching peaks in the simulated spectra, too.
5. We do not find peaks in the MS1 spectrum at *m/z* plus 18.01 (indicating water loss), plus 43.99 (carbon dioxide loss) or minus 17.03 (ammonium adduct); similarly, we do not find molecular formulas in PubChem or ChemSpider that correspond to the novel molecular formula plus H_2_O, plus CO_2_ or minus NH_3_. To this end, we argue that the reported molecular formula indeed corresponds to the protonated molecule, not an adduct or fragment.
6. The spectrum matches to multiple NIST17 library spectra with different *m/z*, all of which are phosphatidylcholines. The top 18 matches have a cosine score above 0.9 and share a set of characteristic peaks which match to the query spectrum: See Fig. 3b for the best hit, and Supplementary Fig. 11 and 12 for additional hits. For the set of shared peaks, the molecular formula annotations of the NIST reference spectrum (as provided by NIST) are identical to those of the SIRIUS fragmentation tree computed for the query compound (Fig. 3e).

Considering the matching NIST reference spectra, we propose that the query compound is a brominated phosphatidylcholine. Marine algae are known producers of halogenated compounds^38^. Moreover, diatoms posess the biosynthetic pathways to produce halogenated lipids^39^. Based on the fragmentation tree analysis and supported by biosynthetical considerations, we propose that the bromine atom is located on the fatty acid tail^40,41^. The putative structure and their mass fragments are shown in Fig. 3d.

Finally, note that there is another novel molecular formula in the diatoms dataset annotated with high confidence, namely C_24_H_49_BrNO_8_P, which differs from the molecular formula C_24_H_47_BrNO_8_P by one degree of unsaturation. The corresponding compounds have *m/z* 590.246 and retention times 523.39 sec and 539.29 sec.

### Running times and stability

In practice, application of Gibbs sampling can be limited by high time demand for burn-in and for sampling a reasonable number of epochs. To avoid this problem, we have used extensive algorithm engineering to reduce running times, as detailed in Section Materials & Methods. Running times were measured on a computer with 40 cores (2x Intel XEON 20 Core E5-2698). We used ten parallel chain, a burn-in of 1,000 epochs and sampling of 2,000 epochs. ZODIAC required between 1 and 14 min per dataset, whereas SIRIUS required between 3 and 42 min per dataset, see Fig. 4. SIRIUS required most time for the dendroides dataset, which contains many high mass compounds. For dendroides, NIST1950 and mice stool, ZODIAC computation did not add much to the total running time whereas for tomato and diatoms, ZODIAC accounts for roughly one-third of the total running time. In all cases, ZODIAC running time is governed by constructing the similarity network of molecular formula candidates, whereas running the Gibbs sampler has a negligible impact. We did not evaluate our optimized Gibbs sampler against a naïve version but from theoretical considerations in Section Materials & Methods, we estimate that the achieved speedup is about 25-fold.

**Fig. 4:**
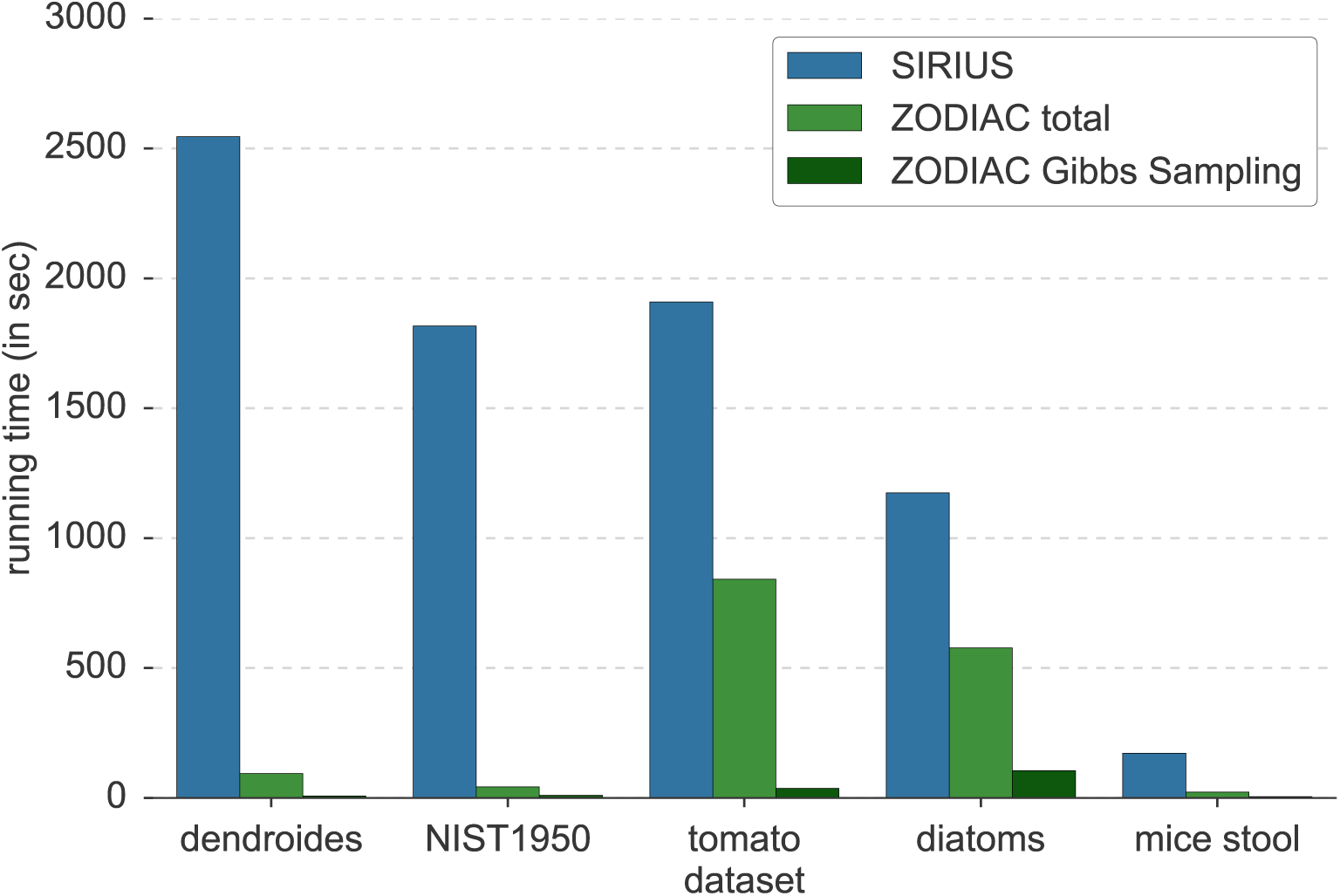
Running time comparison of SIRIUS and ZODIAC on five datasets. We run SIRIUS and ZODIAC on 2× Intel XEON E5-2698 with 40 cores total. “ZODIAC total” running time includes estimation of the edge score distribution, construction of the similarity graph and computation of ZODIAC scores via Gibbs sampling; the later running time is also given separately. ZODIAC requires SIRIUS results as input, and total processing time is SIRIUS time plus ZODIAC time.

In practice, we can speed up the construction of the similarity network, which depends quadratically on the total number of candidates: Here, we used the top 50 candidates for each compound; this conservative approach avoids the exclusion of correct molecular formulas, and also demonstrates the swiftness of our Gibbs sampling method. But running times can easily be reduced by considering fewer candidates, in particular for low mass compounds where SIRIUS usually ranks correct molecular formula much higher. Consequently, ZODIAC can be integrated into existing pipelines without substantial increase in running times.

With regards to stability and required number of epochs, we see that in the beginning both, the total network score and the number of correct molecular formula annotations, are increasing. After 500 to 1,000 epochs the Markov chains reach different local optima. Nevertheless, estimating the most likely candidates from each chain individually results in 96.95% correct molecular formula annotation in all 10 cases. In practice, we run 10 parallel Markov chains to allow for parallelization and to make sampling more robust.

## 4 Conclusion

We have presented ZODIAC, a Gibbs sampling-based approach for assigning molecular formulas in biological samples analyzed by LC-MS/MS. Using ZODIAC, we observed substantial improvements of correct molecular formula annotations, in particular for large compounds; error rates decrease up to 16-fold. Furthermore, the ZODIAC score allows to select the most confident annotations. Different from many other approaches, ZODIAC is not limited to molecular formulas present in any (spectral or structural) databases. We have seen that this is not only of theoretical interest: We confirmed a novel molecular formula discovered by ZODIAC which is, as of today, not contained in PubChem or ChemSpider.

We found that adduct annotations are very important for molecular formula assignment, as it is challenging to deduce this information from isotope pattern and MS/MS data. Hence, high-quality adduct annotations should be established during preprocessing. In contrast, we observed that anchors (library hits) have only a small effect on molecular formula annotations.

Searching an unknown compound with novel molecular formula in a structure database will always result in an incorrect hit, and this will often go unnoticed. In contrast, a metabolite identification workflow which makes use of *de novo* annotation methods facilitates the identification of highly interesting, new metabolites. Here, ZODIAC constitutes a major step in the discovery and structural elucidation of novel metabolites, natural products, and other molecules of biological interest.

## Code availability

ZODIAC has been implemented in the SIRIUS software and is publicly available at https://bio.informatik.uni-jena.de/software/sirius/. The modified version of OpenMS 2.4.0 is available on GitHub (https://github.com/marcus-ludwig/0penMS/tree/release_2.4.0_modifications_Z0DIAC_workflow). A patch file containing changes to OpenMs 2.4.0 is available at https://bio.informatik.uni-jena.de/data/.

## Data availability

Input mzML/mzXML files for the five datasets are available at MassIVE (https://massive.ucsd.edu/), with the following accession numbers: dendroides (MSV000080502), NIST1950 (MSV000081364), tomato (MSV000081463), diatoms (MSV000081731) and mice stool dataset (MSV000079949). SIRIUS and ZODIAC results and a virtual machine to reproduce the data are available from https://bio.informatik.uni-jena.de/data/.

## Acknowledgments

We thank Michael Witting for valuable discussions and Fleming Kretschmer for the fragmentation tree visualization. We gratefully acknowledge financial support by the Deutsche Forschungsgemeinschaft (BO 1910/20) to S.B., K.D., M.F, M.H. and M.L.

## 5 Online Materials & Methods

Throughout this paper, the entities of interest consist of signals detected by mass spectrometry where one or more MS/MS spectra have been recorded by the instrument. It is understood that not all of these signals correspond to compounds in the biological sample; but clearly, only those signals that do correspond are of interest for our analysis. It is also understood that we usually cannot ultimately decide whether a certain signal stems from the protonated molecule [M + H]^+^ or, say, the protonated molecule with a water loss [M−H_2_O + H]^+^ or an ammonia adduct [M + NH_3_ + H]^+^. This is not a problem of our method but rather a general problem of mass spectrometry. For the sake of readability, we will nevertheless use the term “compound” instead of “hypothetical compound”, “feature”, “adduct” or “ion”. In contrast, our methods decide for each compound if it is protonated [M + H]^+^, a sodium adduct [M + Na]^+^ or a potassium adduct [M + K]^+^; in evaluation, compounds that are assigned a wrong adduct are also assigned a wrong molecular formula and, hence, are always counted as misannotations.

### 5.1 Datasets

#### Dendroides

The extract and fractions of *Euphorbia dendroides* plants collected in Corsica were analyzed by HPLC-MS/MS in data dependent acquisition mode on a Orbitrap (LTQ-XL Orbitrap) in positive ionization mode. These same samples^42^ were previously investigated and enabled the isolation and identification of novel diterpene esters, including some endowed with antiviral activities against the chikungunya and the human immunodeficiency virus^43^. These structurally complex and large molecules are characteristic of the Euphorbia genus, and the MS/MS spectra of the isolated diterpene esters were deposited in the GNPS library (CCMSLIB00000840316 to CCMSLIB00000840340).

#### NIST1950

A serial dilution of a plasma methanol extract prepared from the human plasma NIST reference material (SRM 1950)^44^, was analyzed in data dependent acquisition mode by UHPLC-MS/MS on the Q Exactive Orbitrap mass spectrometer in positive ionization mode. Previously, more than 322 compounds were identified by LC-MS in the SRM 1950 reference material^44^. Here, we consider only compounds that were annotated by MS/MS spectral matching with spectra from the NIST17 MS/MS library.

#### Tomato

Fresh tomato seedlings samples (*Solanum lycopersicum*) were extracted in acidic aqueous methanol, offering a wide range of plant metabolites^45^. Samples were analyzed in data dependent acquisition mode by UHPLC-MS/MS on a Q Exactive Orbitrap mass spectrometer in positive ionization mode. Chromatographic separation was performed on mixed-mode C_18_ columns allowing weak anion/cation exchange, that results in the retention of both medium polarity compounds and apolar compounds. We expect a large number of plant metabolite annotations by spectral library search, including phenylpropanoids and terpenoids.

#### Diatoms

This dataset consists of solid phase (PPL, Agilent) extracts of the intra- and exo-metabolomes of a single diatom genus, a major group of marine microalgae. In total, five culture samples of the species *Pseudo-nitzschia subpacifica* (2×), *Pseudo-nitzschia delicatissima* (2×) and *Pseudo-nitzschia multiseries* (1×) were grown in culture from environmental isolates. Cultures included each diatom species and its associated microbiome resulting in a complex and diverse pool of metabolites. We expect the occurrence of compounds containing uncommon elements: Marine microorganisms can contain halogenated organic compounds such as brominated molecules^36^; additionally, the growth medium contained uncommon elements such as silicon and selenium. The samples were analyzed in data dependent acquisition (DDA) mode by UHPLC-MS/MS on a Q Exactive Orbitrap mass spectrometer in positive ionization mode. Metabolomics studies of marine algae are rare, and so, existing libraries are not expected to have many relevant entries, making this an interesting dataset for testing novel compound identification.

#### Mice stool

Quinn *et al.* used this dataset to examine the difference between germ free and colonized mice by metabolomics^46^. In particular, some molecules were only observed in colonized mice, including novel conjugated bile acids and other food derived plant metabolites, which showed the role of the microbiome in their metabolization. Mice stool samples from a microbiome study were analyzed by UHPLC-MS/MS in data dependent acquisition mode mode on a maXis QTOF mass spectrometer in positive ionization mode. Due to technical limitations, the maXis QTOF used for mass spectrometry employs a broader isolation window of (at least 4 *m/z*), and makes identification by computational methods challenging, because isotope peaks can be found in the MS/MS; it also strongly increases the chance of chimeric spectra (fragmentation spectra comprised of fragments from multiple compounds, see below).

### 5.2 Sample preparation and LC-MS/MS analysis

Mass spectrometry data is deposited on MassIVE (https://massive.ucsd.edu/) and MassIVE accession numbers are specified. The exact set of analyzed mzML/mzXML input files is listed in Supplementary Table 7.

#### Dendroides dataset

##### Sample Preparation

The latex of *Euphorbia dendroides* was collected and an ethyl acetate extract was prepared as described by Esposito *et al.*^42^. The extract was then fractionated in 17 fractions that were subjected to mass spectrometry analysis. Subsequent purification led to the isolation and structural isolation of thirteen diterpene esters characterized by extensive NMR spectroscopy and X-ray crystallography diffraction analysis.

##### Mass Spectrometry Analysis

Mass spectra were acquired between *m/z* 150 and *m/z* 1,000. In the full scan mode, full width at half maximum mass resolution of the Orbitrap mass analyzer was fixed at 30,000 for MS spectra and at 15,000 for MS2 spectra. Data-dependent MS^*n*^ mode was used to monitor 1 to 3 most intense ions with an exclusion duration of 40 sec after 8 repetitions. Instrumental parameters were set as follows: source voltage: 5 kV, lens 1 voltage: −15 V, capillary temperature: 275 °C, gate lens voltage: −35 V, capillary voltage: 25V, tube lens voltage: 65 V. The CID parameters were set as follow: CE at 30% of the maximum and an activation time of 30 ms. HPLC was performed with an HPLC Ultimate 3000 system (Dionex, Voisins-le-Bretonneux, France) consisting of a degasser, a quaternary pump, an autosampler, a column oven, and a photodiode array detector. Separation was achieved using an octadecyl column (Sunfire, 150 mm × 2.1 × mm 3.5 μm; Waters, Guyancourt, France), equipped with a guard column. Column oven temperature was set at 25 °C. Elution was conducted with a mobile phase consisting of water + 0.1% formic acid (A) and MeCN + 0.1% formic acid (B), following the gradient 5 to 95% B in 40 min, then maintaining 100% B for 10 min at a flow rate of 250 μL/min. Injection volume was fixed at 10 μL.

##### Data Availability

The mass spectrometry data were deposited on MassIVE (MSV000080502).

#### NIST1950 dataset

##### Sample Preparation

The NIST SRM-1950 human plasma samples were prepared and extracted with 80% ethanol as proposed in the SRM 1950 paper^44^.

##### Mass Spectrometry Analysis

The SRM1950 samples were analyzed using an ultra-high performance liquid chromatography system (Vanquish, Thermo) coupled to an Orbitrap mass spectrometer (Q Exactive, Thermo) fitted with a heated electrospray ionization (HESI-II, Thermo) probe. Chromatographic separation was accomplished using a Kinetex C18 1.3 μm, 100 Å, 2.1 mm × 50 mm column fitted with a C18 guard cartridge (Phenomenex) with a flow rate of 0.5 mL/min. 5 μL of extract was injected per sample/QC. The column compartment and autosampler were held at 40 °C and 4 °C respectively throughout all runs. Mobile phase composition was: A, LC-MS grade water with 0.1% formic acid (v/v) and B, LC-MS grade acetonitrile with 0.1% formic acid (v/v). The chromatographic elution gradient was: 0.0 - 2.0 min, 5% B; 2.0 - 10.0 min, 100% B; 10.0 − 12.0 min, 100% B; 12..0 − 12.5 min, 5% B; and 12.5 − 14.5 min, 5% B. Heated electrospray ionization parameters were: spray voltage, 3.5 kV; capillary temperature, 268.0 ° C; sheath gas flow rate, 52.0 (arb. units); auxiliary gas flow rate, 14.0 (arb. units); auxiliary gas heater temperature, 433.0 °C; and S-lens RF, 60 (arb. units). MS data was acquired in positive mode using a data dependent method with a resolution of 35,000 in MS1 and a resolution of 17,000 in MS2. An MS1 scan from 100-1,500 *m/z* was followed by an MS2 scan, using collision induced dissociation, of the five most abundant ions from the prior MS1 scan.

##### Data Availability

The mass spectrometry data were deposited on MassIVE (MSV000081364).

#### Tomato dataset

##### Sample Preparation

Tomato (*Solanum lycopersicum* var. Better Boy) seeds were surface sterilized in 1.0% (v/v) sodium hypochlorite for 15 min with moderate agitation, rinsed with 20 volumes of sterile distilled water five times, and sown in pots containing peat. Seeds were incubated in a growth chamber with 16h light/8h dark photoperiod and 25 ±2 °C temperature. Two-weeks after germination, roots were washed, dried and whole seedlings were frozen in liquid nitrogen and stored at −80C until further extraction. Three seedlings were pooled into 2 mL tubes and 1.2 mL of acidified aqueous methanol was added (75% methanol (v/v), 24.9% water (v/v), and 0.1% formic acid (v/v)) to obtain a wide range of plant metabolites^45^. The samples were then homogenized in a tissue-lyser (QIAGEN) at 25 Hz for 5 min, and then centrifuged at 15,000 rpm for 15 min. The supernatant was collected in 96 well plates and dried with a vacuum centrifuge. The samples were resuspended in 130 μL of 7/3 MeOH/H2O containing 0.2 μM of amitriptyline (*m/z* 278.189, 542 sec) as an internal standard. After the plates centrifugation at 2,000 rpm for 15 min at 4C, 100 μL of samples were transferred into a new 96 well plate for mass spectrometry analysis.

##### Mass Spectrometry Analysis

Samples were analyzed with an ultra high performance liquid chromatography device (Vanquish, Thermo Scientific) coupled to a quadrupole-Orbitrap mass spectrometer (Q Exactive, Thermo Scientific). Chromatographic separation was performed in mixed-mode on a Scherzo SM-C18 (Imtakt, Torrance, USA) column allowing weak anion/cation exchange (250 × 2 mm, 3 μm) with a guard cartridge (Imtakt). The column was maintained at 40 °C. The mobile phases used were 0.1% formic acid in water (A) and 0.1% formic acid in acetonitrile (B), and flow rate was set to 0.5 mL/min. Chromatographic elution method was set as follows: 0.00 −5.00 min, isocratic 2% B; 5.00 to 8.00 min, gradient 2% to 50% B; 8.00 to 13.00 min, gradient 50% to 100% B; 13.00 −14.00 min, isocratic 100% B; 14.00 − 14.10 min, 100% to 2% B;14.10 to 18.00 min, isocratic 2% B. The injection volume was set to 10 μL. The analyses were performed in electrospray ionization, operating either in positive or in negative ionization mode with a heated electrospray ionization source. In positive ionization mode, the following source parameters were used: spray voltage, +3,000 V; heater temperature, 370 °C; capillary temperature, 350 °C; S-lens RF, 55 (arb. units); sheath gas flow rate, 55 (arb. units); and auxiliary gas flow rate, 20 (arb. units). In negative ionization mode, the following source parameters were used: −3000.0 V; heater temperature, 375 °C; capillary temperature, 350 °C; S-lens RF, 55 (arb. units); sheath gas flow rate, 55 (arb. units); and auxiliary gas flow rate, 20 (arb. units). The MS1 scans were acquired at a resolution of 35,000 (at *m/z* 200) for the 100−1,500 *m/z* range, and the MS2 scans at a resolution of 17,500 from 0.48 to 16.0 min. The automatic gain control (AGC) target and maximum injection time were set at 5 × 105 and 150 ms for MS1 and MS2 scans. Up to four MS2 scans in data-dependent mode were acquired for most abundant ions per duty cycle, with a starting value of *m/z* 70. Higher-energy collision induced dissociation was performed with a normalized collision energy of 20, 35, 50 eV. The apex trigger mode was used (2−7 sec) and the isotopes were excluded. The exclusion parameter was for the data dependent parameter was set to 9 sec.

##### Data Availability

The mass spectrometry data were deposited on MassIVE (MSV000081463).

#### Diatoms dataset

##### Sample Preparation

Diatoms of the genus *Pseudo-nitzschia*, were isolated from waters of the Scripps Pier and their associated microbiome were cultured in filtered Natural Seawater Media, supplemented with inorganic nutrients (NaNO_3_, NaH_2_PO_4_, Na_2_SiO_3_), vitamins and AQUIL trace metals^47^. Cultures were harvested in stationary growth phase and media and intracellular metabolites were solid phase extracted using Bond Elute PPL resin (Agilent) to retain a wide range of non-polar to polar molecules^48,49^.

##### Mass Spectrometry Analysis

The methanol extracts were analyzed by UHPLC-MS/MS on a Q Exactive Orbitrap mass spectrometer in positive data dependent acquisition mode^49^. In short, dried samples were re-dissolved in 100 μL methanol/formic acid (99:1, Fisher Scientific, San Diego, USA) of which 10 μL were injected into a Vanquish UHPLC system coupled to a Q-Exactive Orbitrap mass spectrometer (Thermo Fisher Scientific, Bremen, Germany). For UHPLC separation, a reversed phase C18 porous core column (Kinetex C18, 150 × 2 mm, 1.8 μm particle size, 100 A pore size, Phenomenex, Torrance, USA) was used. As mobile phase A H2O + 0.1 formic acid (FA) and solvent B acetonitrile (ACN) + 0.1% FA was used. The flow rate was set to 0.5 mL/min and after injection, compounds were eluted with a gradient from 0−0.5 min, 5% B, 0.5−8 min 5−50% B, 8−10 min 50−99 B, followed by a 2 min washout phase at 99% B and a 3 min re-equilibration phase at 5% B. ESI parameters were set to 52 L/min sheath gas flow, 14 L/min auxiliary gas flow, 0 L/min sweep gas flow and 400 °C auxiliary gas temperature. The spray voltage was 3.5 kV and the inlet capillary 320 °C. S-lens voltage was 50 V. MS scan range was 150-1,500 *m/z* with a resolution at *m/z* 200 (R_*m/z*200_) of 70,000 with one micro-scan. The maximum ion injection time was set to 100 ms with an AGC target of 1.0E6. Up to 5 MS/MS spectra per MS1 survey scan were recorded DDA mode with R_*m/z*200_ of 17,500 and one micro-scan. The maximum ion injection time for MS/MS scans was 100 ms with an AGC target of 3.0E5 ions and minimum 5% C-trap filling. The precursor isolation width was set to *m/z* 1. Normalized collision energy was set to a stepwise increase from 20 to 30 to 40% with default charge state *z* = 1. MS/MS scans were triggered at the apex of chromatographic peaks within 2 to 15 s from their first occurrence. Dynamic exclusion of precursors was set to a duration 5 s and precursors with unassigned charge states as well as isotope peaks were excluded from MS/MS acquisition.

##### Data Availability

The mass spectrometry data were deposited on MassIVE (MSV000081731).

#### Mice stool dataset

##### Sample Preparation

The mice stool samples were obtained and extracted in 70% ethanol as described by Quinn *et al.*^46^.

##### Mass Spectrometry Analysis

The samples were analyzed with an ultra high performance liquid Chromatography device (UltraMate 3000 Dionex, Fisher Scientific, Waltham, MA USA) coupled with a Bruker Daltonics MaXis qTOF mass spectrometer (Bruker, Billerica, MA USA) as described in ref. ^46^. In brief, the metabolites were separated using a Kinetex 2.6 μm C18 (30 × 2.10 mm) UHPLC column fitted with a guard column. The isolation width was dependent on *m/z* value, with a 4 *m/z* isolation for 50 *m/z* to 8 *m/z* at 1,000 or higher. Lower isolation width results in a drop in sensitivity.

##### Data Availability

The mass spectrometry data were deposited on MassIVE (MSV000079949).

### 5.3 Preprocessing

To ensure reproducibility, we provide a virtual machine comprising all steps of the preprocessing. The virtual machine incorporates the OpenMS sources, executables and parameter files and all data processing scripts written in Java and Python. The workflow described below is visualized in Fig. 7.

#### OpenMS

We used OpenMS 2.4.0^34^ to process the mzML/mzXML files. We performed minor modifications on the OpenMS source code by removing 12 lines and adding 78 lines. This allowed to detect more isotope peaks, to match MS/MS to MS1 features based on the actual isolation window and to add functionality to the SIRIUSAdapter to directly output SIRIUS file format including retention time information. These numbers are based on the patch file, see Code availability, and include lines with comments and blank lines; the modification of a line corresponds to the removal and insertion of a new line.

Feature finding and clustering of isotopic mass traces was performed using the FeatureFinder-Metabo module. Next, adducts were detected with the MetaboliteAdductDecharger module. Finally, spectra were exported to the SIRIUS specific format using the SIRIUSAdapter module. OpenMS parameter files are provided as part of the virtual machine. Parameters were chosen by manual inspection; in particular, we used a small noise intensity threshold to increase chances that isotope peaks of a compound are picked.

#### Discarding features and MS/MS spectra

We excluded *m/z* features which eluted over a very long time during chromatography and did not produce desired mass traces in a limited time window, as such traces are considered chemical noise. To do so, we binned MS1 peaks with a bin size of 0.006 *m/z*. Each MS1 was normalized by the most intense peak. Each peak was counted if its relative intensity was 0.01 or higher. If a *m/z* bin contained peaks of more than 20% of MS1 the *m/z* was considered chemical noise. Because these spurious chemical noise features have rather high mass deviation we removed all MS1 features within 30ppm.

Next, we performed blank removal using blank samples from the corresponding datasets. Features within 15 ppm and 20 s of a blank feature were removed if intensities were lower than 2-fold of the blank feature intensity. We did not perform blank removal on the mice stool dataset, because this resulted in a low number of remaining compounds.

Third, we removed features from the beginning and end of the chromatography run and features with low relative or absolute intensity; and we removed MS/MS spectra which could not be assigned to an MS1 feature, MS/MS of a precursor peak with low absolute or relative intensity, and chimeric MS/MS. See Table 2 for dataset-specific parameter values. Chimeric spectra contain fragments of multiple precursor ions; we detected chimeric spectra as follows: All peaks within the isolation window, excluding isotope peaks, were considered to contribute their intensity to the measured MS/MS. We estimated the relative intensity that the target precursor ion contributes to the MS/MS; if the target precursor ion contributed to less than 50% of the MS/MS intensity or if a second precursor ion contributed more than 33% of the target precursor ion intensity, the MS/MS was marked as chimeric and excluded. The Isolation window width for the Orbitrap mass spectrometer used for the dendroides, NIST1950, tomato and diatoms is 1 Da; for the mice stool dataset analyzed on a QTOF mass spectrometer an isolation window of 3 Da width and shifted by 1 Da to the right, centered at the +1 isotope peak, was assumed.

#### Filtering MS/MS spectra

In each MS/MS spectrum, we filtered peaks using an intensity threshold of two times the median noise intensity, see Table 2. The median noise intensity of a dataset was estimated from peaks which had no molecular formula decomposition within a 40 ppm window considering elements C, H, N, O, and P plus those elements predicted from the isotope pattern, see below. Isotope peaks were removed from MS/MS spectra of the mice stool dataset.

The SIRIUSAdapter OpenMS module combines MS/MS which are associated with the same MS1 feature. In addition, complete linkage hierarchical clustering was conducted to merge features over different LC-MS/MS runs. Features were merged using 15 ppm mass accuracy and a 15 sec retention time window. Features with different adduct annotations or features from the same run were not merged. Feature similarity was computed by the cosine product of the MS/MS (see below), and the similarity threshold for clustering was set to 0.8. When multiple features were merged into a single one, where each feature has an assigned isotope pattern, then the isotope pattern with the highest number of isotope peaks was kept. In case multiple isotope patterns had the same number of isotope peaks, the one with the most intense monoisotopic peak was kept. After merging, features were discarded if the summed MS/MS intensity was below a threshold, see Table 2. Features with precursor mass above 850 Da are discarded: Whereas ZODIAC is clearly capable of processing such features, we found that there are no spectral library hits above this mass that can be used for evaluation, see below. Only 2.72% of features across all datasets have *m/z* above 850 Da, so excluding these cannot have substantial impact on result statistics.

#### Extending isotope patterns

OpenMS often misses low-intensity isotope peaks. To recover those peaks, we post-processed OpenMS results as follows: For each isotope pattern detected by OpenMS, we try to extend it using isotope peaks from the corresponding MS1 spectra chosen by OpenMS. Isotope pattern peaks were picked using the SIRIUS 4 isotope pattern picking subroutine. If an additional isotope peak is present in at least 66% of the corresponding MS1, the peak was added to the isotope pattern. Subsequently, features with less than two isotope peaks are discarded.

#### Discarding low-quality merged MS/MS spectra

Even when considering all MS/MS spectra for some features, we sometimes have insufficient information for both spectral library search and molecular formula annotation; to this end, such “low-quality features” were discarded. A feature is discarded if it produces less than 5 fragment peaks, estimated after merging peaks within 10 ppm or 0.0025 *m/z* from all corresponding MS/MS spectra; and if no fragmentation tree in the top 50 candidate list can explain at least 5 peaks accounting for at least 80% of total spectrum intensity, see SIRIUS analysis below. Filtering “low quality” features decreased the number of features for dendroides from 1,078 to 784, for NIST1950 from 568 to 400, for tomato from 3,583 to 2,584, for diatoms from 3,227 to 2,075 and for mice stool from 577 to 377.

For brevity, we will refer to the features detected by OpenMS as *compounds*, see above.

### 5.4 SIRIUS analysis and establishing a ground truth

SIRIUS 4 was run with the default alphabet of elements CHNO, and at most 5 phosphorus atoms; automatic element detection from the isotope pattern^50^ was enabled for sulfur, chlorine, bromine, boron, and selenium. For the diatoms dataset, we added silicon to the set of auto-detectable elements. For the dendroides, NIST1950, and tomato datasets we used 15 ppm maximum mass deviation for SIRIUS; for diatoms and mice stool datasets we used 10 ppm. Isotope patterns were not used to filter molecular formula candidates before computing fragmentation trees.

– If OpenMS provided an ionization adduct type (such as protonation, sodium adduct, potassium adduct) for a compound, only this ionization was used. We export the 50 best-scoring molecular formula candidates from SIRIUS.
– In cases where no ionization adduct type was provided by OpenMS, we selected one or more adducts from [M+ H]^+^, [M+ Na]^+^, and [M+ K] ^+^ by searching for characteristic mass differences, using the MS1 that contained the most intense peak of the precursor ion. Peaks below 5% relative intensity were discarded for this decision. For each compound, we export the 50 best-scoring molecular formula candidates; we simultaneously ensure that for each considered ionization adduct type, at least 10 candidates are considered.

#### We will refer to this candidate list as the *top 50*

To evaluate the performance of SIRIUS and ZODIAC, we had to annotate a subset of compounds with “correct” molecular formulas, to serve as our ground truth. For this, we combined manual annotation and spectral library search, as follows: For the dendroides dataset, spectral library hits were obtained for the isolated molecules that had their reference MS/MS spectra added to the GNPS library. We used molecular networking and spectral library search in analog mode^1^, along with a set of known typical biotransformation, to annotate related diterpene esters. They differ mainly by their acylation degree, and the nature of acyl residues on the diterpene backbone. This resulted in 201 compounds being annotated with molecular formulas by manual analysis of the data, see Supplementary Table 5.

For the remaining datasets, we performed spectral library searches against multiple libraries, but did not add manual annotations. We searched compounds in a spectral library combining GNPS^1^, MassBank^4^, NIST17 database (National Institute of Standards and Technology, v17) and “MassHunter Forensics/Toxicology PCDL library” (Agilent Technologies, Inc.)^10^. We compute a similarity score assuming peaks as Gaussians, with the centroided peaks’ *m/z* as the mean and the standard deviation being the maximum of a relative mass error of 20 ppm and an absolute mass error of 0.005 *m/z*. Precursor ion masses are permitted to differ by 10 ppm or 0.0025 *m/z* at maximum. Only library hits with a similarity score of 0.7 or higher and with at least 6 shared peaks are considered as being valid. We compute the score as the mean of the cosine score of the sample spectrum and the cosine score of the mirrored spectrum; to mirror a spectrum with precursor mass *M*, we replace peak *m/z* value m by *M − m*. This resulted in 94 annotated compound for NIST1950, 271 for tomato, 93 for diatoms and 44 for mice stool, see Supplementary Table 6.

We evaluate SIRIUS and ZODIAC against these “ground truth” molecular formulas, but we stress that beside the molecules that were isolated in *Euphorbia dendroides* samples and correspond to level 1 of the Metabolomics Standard Initiative ranking system, not all of these are necessarily correct. In particular, we refrain from ranking these according to the Metabolomics Standard Initiative ranking system, where level 4 corresponds to an “unequivocal molecular formula”. An evaluation is nevertheless meaningful because we expect only few errors on the molecular formula assignment level.

In few cases, the correct molecular formula was not ranked in the top 50 SIRIUS candidates; we also dropped these from our evaluation, as it is not possible that ZODIAC can find the correct molecular formula in our evaluation. We discarded four compounds for dendroides, zero for NIST1950, two for tomato, zero for diatoms and one compound for mice stool because of this criterion.

See Supplementary Table 1 for details, and see Fig. 6 for the mass distribution of the “ground truth” compounds. Compounds in the dendroides dataset with reference annotations have high mass, and 75% of all reference annotations have an *m/z* of 605 or higher. The NIST1950 dataset resulted in library hits over a broad range of *m/z* values. The diatoms library hits have a median *m/z* of 301 but the sample itself is highly complex, as described above. Only few compounds remain in the mice stool dataset after filtering chimeric and low quality compounds, see above.

### 5.5 Posterior probability of an assignment

We use a probabilistic view on the molecular formula assignment problem^27^: For each hypothetical compound in the LC-MS run, we are given data such as an isotope pattern and a fragmentation pattern. This allows us to determine, for each compound *c* ∈ 𝒞, a set of candidate molecular formulas that may explain the observed data. Let *V* be the set of all molecular formula candidates, such that *V* (*c*) ⊆ *V* is the subset of molecular formulas for compound *c* ∈ 𝒞. It is possible that different compounds share an identical molecular formula explanation, but we ignore this in our presentation, solely for the sake of readability. An *assignment* is a mapping **𝔞**: 𝒞 → *V* where **𝔞** (*c*) ∈ *V* (*c*) is the molecular formula assigned to compound *c*. The posterior probability of an assignment **𝔞** is

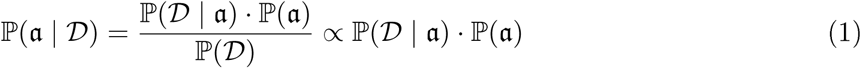

where 𝒟 is the observed data. We use the terms “prior probability”, “likelihood” and “posterior probability” according to this Bayesian point of view. Let 𝒟 (*c*) be the observed data for compound *c* ∈ 𝒞, that is, the isotope pattern and fragmentation pattern of *c*. We assume that the likelihoods of molecular formulas for different compounds are independent, and that the likelihood of any compound *c* only depends on its data 𝒟 (*c*); so,

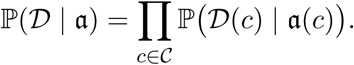

Next, we define the prior probability of an assignment as the product of priors for pairs of compounds:

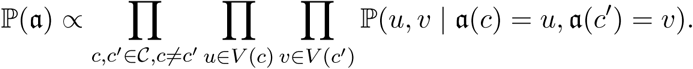

Here, ℙ (*u, v* | “true”) is the prior probability that two compounds with molecular formulas *u, v* co-occur in the dataset; analogously, ℙ (*u, v* | “false”) if *u, v* do not co-occur. To simplify our calculations, we introduce a mapping **𝔠**:*V* → *C* that maps any molecular formula to the compound it belongs to: **𝔠** (*v*) = *c* for all *v* ∈ *V* (*c*), for *c* ∈ *C*. Note that **𝔠** (**𝔞** (*c*)) = *c* for all *c* ∈ 𝒞. Now,

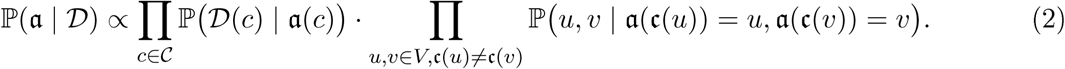

Different from Rogers *et al.*^27^, we are able to formulate the posterior probability of an assignment in closed form. A natural question is if we can find a maximum a posteriori estimate for (2); unfortunately, we will see that this is not easy, as the underlying computational problem is NP-complete. Another natural question is to sample from the posterior distribution; this will be addressed below.

### 5.6 Graph-theoretical formulation

We now give a graph-theoretical formulation of the problem; this will allow us to establish its computational complexity, but also to come up with a more efficient algorithm. Let *V*, the molecular formula candidates, be the nodes of an undirected graph *G* = (*V, E*) with edge set 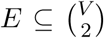. We will write *uv* as shorthand for a tuple 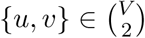. We use **𝔠**: *V →* 𝒞 as a *node coloring* with color set 𝒞. Now, an *assignment* is a subset *A ⊆ V* such that each color from 𝒞 appears exactly once; in this case, *A* is also called *multicolored*. Using the notation of the previous section, we have *A* = **𝔞** (𝒞); recall that **𝔠** (**𝔞** (*c*)) = *c* for all *c* ∈ 𝒞. Let *w*: *V* ∪ *E* → ℝ be *weights* for all nodes and edges of the graph. The *weight* of the assignment *A* is

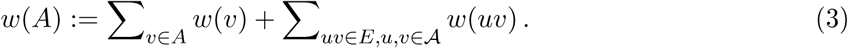

This corresponds to the node plus edge weights of a node-induced subgraph of *G*, for node set *A* ⊆ *V*.

We consider the following optimization problem:

#### Maximum Multicolored Subgraph problem

We are given a graph *G* = (*V, E*), a node coloring 𝔠 : *V* → *𝒞* and weights *w* : *V ∪ E* → ℝ. We search for an assignment *A* ⊆ *V* of maximum weight, that is, a node-induced multicolored subgraph of maximum weight.

How does this problem correspond to our probabilistic problem from the previous section? Setting *E* = *E*\*: = {*uv* | *u*; *v* ∈ *V*; 𝔠(*u*) ≠ 𝔠(*v*)} (the set of all node pairs with different colors) and

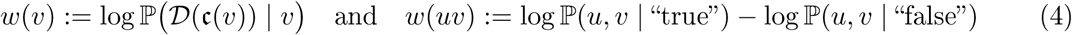

we can show that these problems are in fact equivalent: We have log ℙ (𝔞 | 𝒟) = *w*(𝔞(𝒞)) + *α* for some constant *α* ∈ ℝ. Here, we assumed that *E* = *E*\* contains all possible edges; we call (*V, E*\*) a *complete assignment graph*. But we can encode any edge set *E* ⊊ *E*\* using zero edge weight for all *e* ∉ *E*, so both problems are equivalent. Hence, it is natural to ask for an optimal solution of the problem, which would correspond to a maximum a posteriori estimator.

### 5.7 Complexity of the problem

For the decision version, we ask if there is an assignment with weight above some threshold *τ* ∈ ℝ. In its simplest form, all edges have weight one and all nodes have weight zero, *w*|_*E*_ ≡ 1 and *w*|_*V*_ ≡ 0.

#### Lemma 1.

*The* MULTICOLORED SUBGRAPH *problem is NP-complete, even for unit edge weights and zero node weights.*

*Proof.* It is clear that the MULTICOLORED SUBGRAPH problem is in NP. We show that the problem is NP-hard by reduction from CLIQUE^51^: Let *G* = (*V, E*) be an undirected, simple graph, is there a clique of size *k* in *G*? Clearly, *k* ≤ *n*: = |*V* |.

We construct a graph 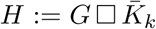 as the Cartesian graph product of *G* and the empty graph 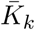 with *k* nodes and no edges: That is, for every node *v* ∈ *V* we generate *k* copies (*v*, 1), …, (*v, k*) in *H*, and there is an edge {(*u, i*), (*v, j*)} with *i* ≠ *j* in *H* if and only if there is an edge *uv* in *G*. Now, *k* ≤ *n* implies that *H* contains at most *n*^2^ nodes. We define node colors 1, …, *k* such that *c* (*v, i*) = *i* for *v* ∈ *V* and 1 ≤ *i* ≤ *k*. We assign zero node weights and unit edge weights for all nodes and edges in *H*. Now, any assignment in *H* corresponds to a *k*-node induced subgraph in *G*, and the weight of the assignment equals the number of edges in the node-induced subgraph; to this end, an assignment of weight 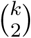 would correspond to a *k*-clique in *G*.

The Multicolored Subgraph problem is a generalization of the Multicolored Clique problem; to this end, Lemma 1 can also be inferred from the complexity of Multicolored Clique, which is W[1]-hard^52^. Assuming zero node and unit edge weights, the above construction implies that for any *ϵ* > 0, there is no polynomial time algorithm that approximates the maximum assignment weight to within a factor better than *O*(*n*^1−*ϵ*^), unless P = NP^53^. Furthermore, finding an assignment of weight *k* cannot be done in time *n*^*o*(*k*)^, unless the exponential time hypothesis fails^54,55^. Finally, we noted above that we can encode an arbitrary edge set *E* ⊊ *E*\* using zero edge weight for all *e* ∉ *E*, so:

#### Corollary 1

*The* MULTICOLORED SUBGRAPH *problem is NP-complete, even for a complete assignment graph, binary edge weights and zero node weights.*

Finally, we consider two problem variants: First, we may allow that some colors from 𝒞 are absent from *A*; in this case, *A* is called *colorful*. We can encode this variant in the original problem, by adding a dummy node for each color which is connected to no other node. Second, we may assume that only edges carry weight. We can encode the MULTICOLORED SUBGRAPH problem in this variant, by adding a dummy color for each color and a dummy node for each node, such that if a node has a certain color, then the dummy node has the corresponding dummy color. We connect each node to its dummy node, and transfer the weight of the node to the corresponding edge. Hence, our complexity results also hold for these variants.

On the algorithmic side, it is easy to see that the MULTICOLORED SUBGRAPH problem can be solved by a simple Integer Linear Program (one variable per edge and one variable per color). We omit the straightforward technical details. We will not proceed in this direction, as this approach results in a single optimal solution, whereas we want to consider suboptimal solutions and marginal probabilities, which allow us to judge our individual confidence when assigning molecular formulas to compounds.

### 5.8 Likelihoods, prior probabilities and graph topology

The likelihood ℙ (𝒟(𝔠 (*v*))|*v*) of a molecular formula candidate *v* can be computed from the posterior probability of the fragmentation tree and the isotope pattern analysis as estimated by SIRIUS 4.0^10,24^. For the Gibbs sampler, we treat these probabilities as likelihoods, although the analysis SIRIUS 4.0 also integrates certain priors^24^. To avoid proliferating running times, we usually limit further computations to the, say, 50 best-scoring molecular formulas for each compound. For each compound, we also introduce a node representing “molecular formula not identified” which receives likelihood from the remaining molecular formulas, and is not connected to any other nodes.

Furthermore, we assume that some compounds were identified by searching in a library of tandem MS spectra, plus potentially by comparison of retention times. We refer to these compounds and the corresponding molecular formulas as “*anchors*”. Such library search results can also be wrong, so we do not exclude other molecular formula explanations, but rather give a bonus to the likelihood of the identified molecular formula. The “quality” of a spectral library hit can, to a certain extend, be evaluated using its score, usually the dot product (cosine score) between query and reference. Hence, the bonus may be dependent on the corresponding library search score. Given the library search score *s*_*l*_ ∈ [0, 1] and a minimum score to consider a library hit *min*_*l*_, we multiply the candidate’s likelihood by

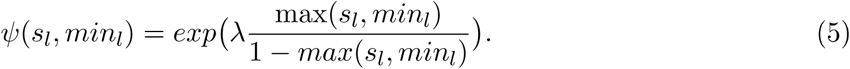

Candidates which disagree with the library hit or without any library hit are scored using *s*_*l*_ = *min*_*l*_. Note, that any “perfect match” with score of 1.0 will be chosen in any case. We remove any other candidate for this compound. We refrain from normalizing the *ψ* to one.

For estimating priors, we will consider similarity of fragmentation patterns^32,33^: More precisely, we use similarity between fragmentation trees that were computed by SIRIUS in the previous step. For each pair of compounds, we have to compare up to 50 times 50 fragmentation trees: For swift computations, we refrain from using fragmentation tree alignments^56^ but instead, simply count the number of common fragments and precursor (root) losses in the two trees^56^. Evaluations indicate that this method, while performing worse than fragmentation tree alignments, is still able to detect structural similarity between compounds^56^. When counting common root losses, the empty root loss is ignored. We introduce two modifications to the score from^56^: Let *n*_1_, *n*_2_ be size of the two fragmentation trees, defined by the number of fragments and root losses. Instead of normalizing the number of common fragments plus root losses *s* by the size of the smaller tree min{*n*_1_, *n*_2_}, we use

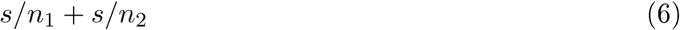

as the normalized score; by this, we slightly penalize large trees, as having common fragments or root losses is more likely against a large than a small tree. But this score favors small trees and, hence, inferior molecular formula candidates. To this end, we use the size of the *largest* fragmentation tree, among all candidate molecular formulas, for the normalization of each compound; this is the maximum number of explainable peaks in the tandem MS data of the compound. Fragments and root losses can be weighted by importance *ι*. The weight of two common fragments or root losses *m*_1_ and *m*_2_ is *ι*(*m*_1_)*ι*(*m*_2_). The weighted size of a tree is

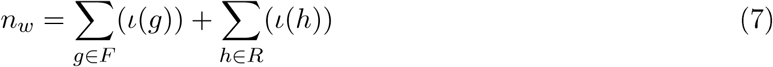

with fragments *F* and root losses *R*. For two molecular formulas *u, v* ∈ *V* we denote the resulting score as *s*(*u, v*).

How can we transform this count into a prior probability? Natural choices include significance estimates such as p-values and posterior error probabilities. We do not have a reasonable model for the score distribution of “true” edges; in fact, it is not know how to clearly distinguish between “true” and “false” edges in such a model. To this end, we resort to a simple prior based on p-value estimation:

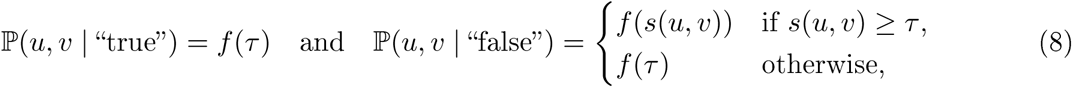

where *τ* ∈ ℝ is a thresholding parameter, and *f*: ℝ → [0, 1] is a monotonically decreasing function. We introduce threshold *τ* because scores below a certain threshold are practically uninformative and should not be considered in our estimations. For *f* (*x*) we estimate the p-value of score *x*, under the null model that scores follow a certain distribution. Note that prior probabilities do in fact depend upon the (mass spectrometry) data.

We now assign node and edge weights according to (4). Clearly, many of these edges have zero weight and can be removed from the graph. To avoid that nodes are isolated, we want to keep some edges incident to any node. This can be formulated by *individual thresholds τ*_*c*_ ∈ ℝ for each color *c* ∈ 𝒞 and, for an edge *uv*, edge weight

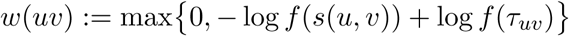

for threshold *τ*_*uv*_: = min {*τc*(*u*), *τc*(*v*)} This will change the weight of any assignment by an additive constant and, hence, posterior probability by a multiplicative constant.

### 5.9 (Faster) Gibbs sampling

We say that a node *v* is *active* in an assignment *A* if *v* ∈*A*, and that an edge *uv* is *active* if both *u* ∈ *A* and *v* ∈ *A*; then, the weight of an assignment is the sum of weights of all active nodes and edges.

Gibbs sampling is a Markov chain Monte Carlo algorithm for obtaining a sequence of observations approximated from a multivariate probability distribution^57^. Sampling assignments according to (2) can be seen as an archetype application of a Gibbs sampler: We start with some assignment, such as the highest likelihood node (molecular formula) for each compound (color). Each epoch of the Gibbs sampler consists of | 𝒞 | *steps*, where we iterate over all colors *c* ∈ 𝒞 in random order: We update the active node with color *c* by drawing a node with color *c* according to its posterior probability, conditional the current assignment of all nodes with color different from *c*. At the end of the epoch we output the current assignment, and repeat until we have reached a sufficient number of samples. This generates a Markov chain of samples converging to the posterior probability distribution of assignments. In practice, we discard samples from the beginning of the chain (*burn-in period*), and to avoid correlation between nearby samples, we output only every, say, 10^th^ sample.

Assume that *u* ∈ 𝒜 with color *c*: = **𝔠** (*u*) is to be (potentially) replaced by a new node *v* with the same color. The probability of *v* ∈ *V* (*c*), conditional all other nodes *z* ∈ *A* with **𝔠** (*z*) ≠ *c*, can naïvely be computed as

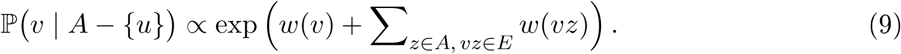

Computing all conditional probabilities for drawing a node *v*, requires time proportional to the sum of node degrees for all nodes from *V* (*c*). That means running time for one step is of order *O*(|*V* (*c*) | · |*V|*) and, hence, *Θ*(|*V*| ^2^) for certain graph families.

To apply Gibbs sampling in practice, the critical point is to quickly reach a large number of samples, so that probability estimates become reliable. To further decrease running time, we assume that we have, at any step, knowledge about all (log) conditional probabilities, for all nodes *v* ∈ *V* (*c*) and all colors *c* ∈ 𝒞. We assume that conditional probabilities are not normalized; to sample a new active node, we uniformly draw a random number between zero and the sum of conditional probabilities, over all nodes with this color. To improve the sampling speed, we want to estimate conditional probabilities without performing a full calculation using (9).

#### Lemma 2

*One step of the Gibbs sampler, exchanging some node u by another node v with the same color c*: = **𝔠** (*u*) = **𝔠** (*v*), *can be carried out in O* (|*V* (*c*)| + deg(*u*) + deg(*v*)) *time.*

*Proof.* Let *A* ⊆ *V* be the current assignment with *u* ∈ *A*. We want to choose a new node *v* ∈ *U* from the set of candidate nodes *U*: = *V* (*c*) for color *c*: = **𝔠** (*u*). We know the conditional probabilities ℙ(*v* | 𝒜− {*u*}) for all *v* ∈ *U*; we sum up the conditional probabilities, then uniformly choose a random number between zero and this sum and, finally, use this random number to select one *v* ∈ *U*. This can be carried out in time *O*(|*U* |). If *u* = *v* then we can stop at this point.

Second, we have to estimate conditional probabilities for all nodes *z* ∈ *V*. From (9), we infer that the conditional probability only changes for those nodes *z* where there is a change in the neighborhood *N* (*z*) of *z*, and remains constant for all others. To this end, we iterate over all *z* ∈ *N* (*u*), and decrease the log conditional probability of *z* by *w*(*uz*); then, we iterate over all *z* ∈ *N* (*v*), and increase the log conditional probability of *z* by *w*(*vz*). Finally, for any node *z* ∈ *N* (*u*) ⋃ *N* (*v*), we recompute its conditional probability using the exponential function. This can be carried out in time *O*(deg(*u*) + deg(*v*)); afterward, all conditional probabilities are correct for the new assignment 𝒜 − {*u*} ∪ {*v*}.

Comparing a naïve graph-based implementation of a Gibbs sampler with one that uses Lemma 2, we can estimate that the speedup is of order *Θ*(| *V* (*c*) |).

For the first iteration, we use an arbitrary assignment, then compute all conditional probabilities using (9). The method requires *O*(|*V*| + |*E*|) memory for storing the graph, and *O*(|*V*|) memory for storing (log) conditional probabilities. The probability of a particular molecular formula *v* to be correct, can now be estimated as its marginal probability: that is, the ratio of assignments in the output that contain *v*.

### 5.10 ZODIAC parameters

We use identical parameters for all five datasets, see (5) above: We weight fragments and root losses when comparing fragmentation trees of molecular formula candidates. Here, we use the SIRIUS 4 noise intensity scoring as importance *ι* in (7). The probability that a peak *p* that corresponds to a fragment and root loss is not noise is *ι* = 1 − *par*(*int*(*p*)), where *par* is the Pareto cumulative distribution function with *x*_*min*_ = 0.002, *x*_*median*_ = 0.015 and *int*(*p*) ∈ [0, 1] the relative peak intensity, see ref. ^24^. To establish a threshold on the minimal similarity of fragmentation trees, we decrease score *s* and tree sizes *n*_1_ and *n*_2_ each by 1.0, see (6).

The empirical score distributions resemble a log-normal distribution, see Supplementary Fig. 9, so we use its Cumulative Distribution Function to estimate p-values for (8). For the robust estimation of parameters *μ* and *σ*^2^, we sampled 100,000 non-zero scores for each dataset, and used the median score as parameter *μ* and the median absolute deviation as parameter *σ*^2^. We naturally expect most edges to be false edges and chose score threshold *τ* so that 95% of the non-zero scores are smaller than this threshold. Finally, we use individual thresholds for each compound (color) so that at least 10 molecular formulas of this color are incident to 10 or more edges.

Each molecular formula candidate of some compound receives a score *s*_1_, …, *s*_*n*_, where *s*_*max*_ is the largest score. We transformed SIRIUS scores to probabilities using the softmax function, where *p*_*j*_ = *exp*(*s*_*j*_ − *s*_*max*_) are normalized to sum to one. To adjust for the fact that the correct molecular formula may not be in the top 50, we added a dummy node receiving the combined probability of all unconsidered candidates. Dummy nodes are not connected to any other node. SIRIUS does not report the score of all candidates, as one compound may have tens of thousands of candidates. Hence, we estimated the probability of all unconsidered candidates by multiplying the number of unconsidered candidates with the lowest probability of the top 50 candidates.

#### Finding and scoring ZODIAC anchors

ZODIAC can use (potentially incorrect) spectral library hits as anchors to improve annotations. To find a reasonable number of anchors, we perform spectral library search in analogue mode. Resulting molecular formula annotations are not considered ground truth identifications but are sufficient as anchors. Only those hits were considered that have mass diferences between query and reference corresponding to a frequent biotransformation. We use the following molecular formula mass diferences as valid biotransformations^27,58,59^: C_2_H_2_, C_2_H_2_O, C_2_H_3_NO, C_2_H_3_O_2_, C_2_H_4_, C_2_O_2_, C_3_H_2_O_3_, C_3_H_5_NO, C_3_H_5_NO_2_, C_3_H_5_O, C_4_H_2_N_2_O, C_4_H_3_N_3_, C_4_H_4_O_2_, C_5_H_7_, C_5_H_7_NO, C_5_H_9_NO, CH_2_, CH_2_ON, CH_3_N_2_O, CHO_2_, CO, CO_2_, H_2_, H_2_O, N, NH, NH_2_, NH_3_, O, H_4_O_2_ and H_6_O_3_.

We use identical parameters for all five datasets: When scoring anchors according to (5), we use the *maximum* of the cosine score between the spectrum and the cosine score of the mirrored spectrum as the similarity measure, and *min*_*l*_ = 0.5 as the score threshold parameter and *λ* = 1, 000 as the weighting parameter. For anchors found by spectral library match in analogue mode (that is, non-identical *m/z*), spectral similarity is reduced by 0.1 to account for increased uncertainty.

Searching for anchors as described above resulted in 96 anchors for dendroides, 254 anchors for NIST1950, 749 for tomato, 372 for diatoms and 176 for mice stool. All spectral hits described in the previous section are anchors, too; recall that for dendroides, the ground truth was established manually and those annotations do not serve as anchors.

#### Burn-in and number of Gibbs sampling epochs

We determined a reasonable number of Gibbs sampling iterations using the dendroides dataset. One iteration, also called *epoch*, is defined as one round in which each compound is updated once by choosing a new “active” molecular formula candidate. We run 10 independent Markov chains, see Fig. 5: The total score summed over all active candidate at a specific epoch increases swiftly over the first 500 epochs. Similarly, the number of correct annotations at a specific epoch increases quickly for most Markov chains until the chain seems to stay in a local optimum. We note that this number of correct molecular formula is determined at each epoch whereas ZODIAC scores are computed from the average over many epochs. From this data, we estimated a burn-in of 1,000 epochs and sampling of 2,000 iterations. Larger values increase running times but should never worsen results.

**Fig. 5:**
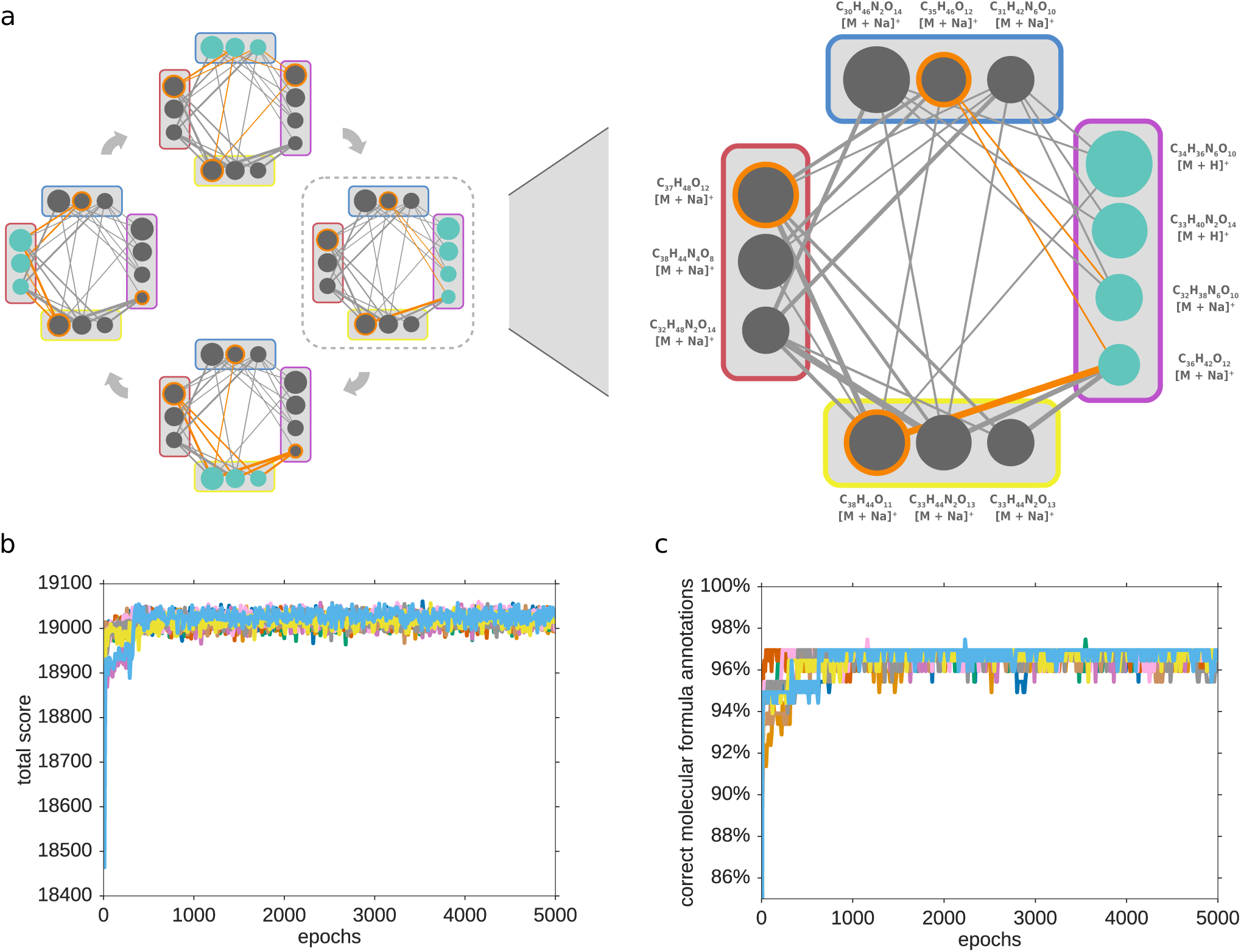
Gibbs sampling. (a) Illustration of the Gibbs sampling process. Left: During each epoch compounds are iterated in random order. For each compound a new active molecular formula candidate is sampled based on prior probabilities and active candidates of all other compounds. Right: Sampling step for one compound. This illustrated sub-network with four compounds is based on the dendroides dataset. Each circle corresponds to a molecular formula candidate. The size depicts the rank estimated by SIRIUS. Orange rings mark active candidates. Edge width depicts fragmentation tree based similarity between candidates. The candidates which are sampled in this step are colored cyan. The bottom figures show the total assignment score of the molecular formula candidate network (b) and identification rate over the course of epochs (c) for Gibbs sampling on the dendroides dataset.

In application, we use 10 Markov chains in parallel, a burn-in of 1,000 epochs, and sample 2,000 epochs; we keep only every 10^th^ sample, resulting in a total of 10 × 200 = 2, 000 samples.

## 6 Supplementary Tables and Figures

**Supplementary Table 1:** Statistics on compounds with annotated ground truth molecular formulas. Given is the number of total compounds, the number of compounds with a ground truth molecular formula and the number which are in the top 50 of SIRIUS ranked candidates. The median *m/z* and 25 and 75 percentile considers only candidates in the top 50.

| dataset | total compounds | ground truth | # in top 50 | 1st quartile $m/z$ | median $m/z$ | 3rd quartile $m/z$ |
| --- | --- | --- | --- | --- | --- | --- |
| dendroides | 791 | 201 | 197 | 605.310 | 705.274 | 759.353 |
| NIST1950 | 568 | 94 | 94 | 286.390 | 373.800 | 477.335 |
| tomato | 2,829 | 271 | 270 | 207.814 | 271.713 | 334.526 |
| diatoms | 2,457 | 93 | 93 | 253.195 | 301.216 | 349.237 |
| mice stool | 390 | 44 | 43 | 373.274 | 454.292 | 516.298 |

**Supplementary Figure 6:**
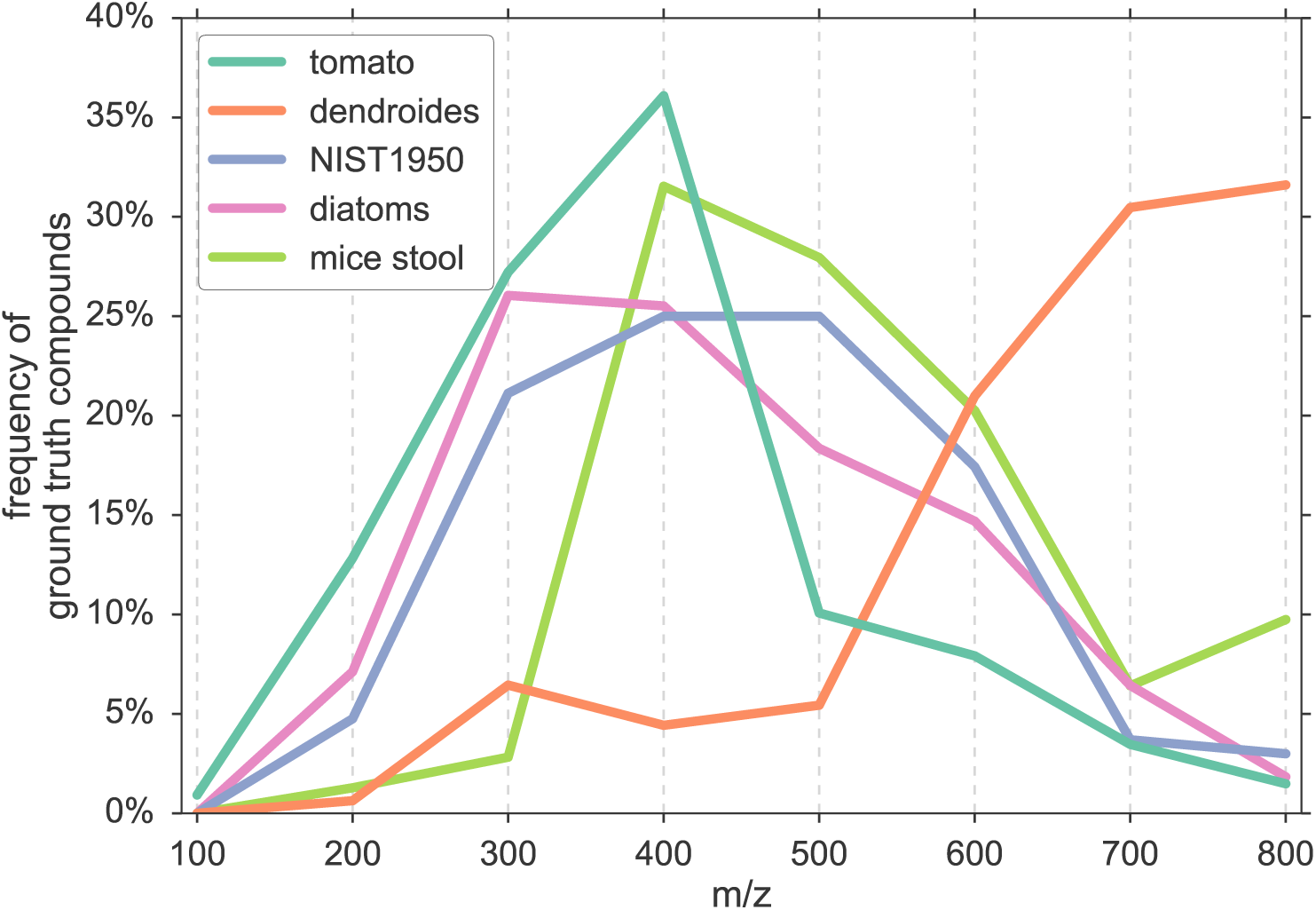
Distribution of compound masses. Distribution of precursor ion *m/z* of the compounds used as ground truth for the evaluation of the molecular formula annotation on the five datasets. Bins of width 100 are centered at 100, 200, …, 800 *m/z*.

**Supplementary Figure 7:**
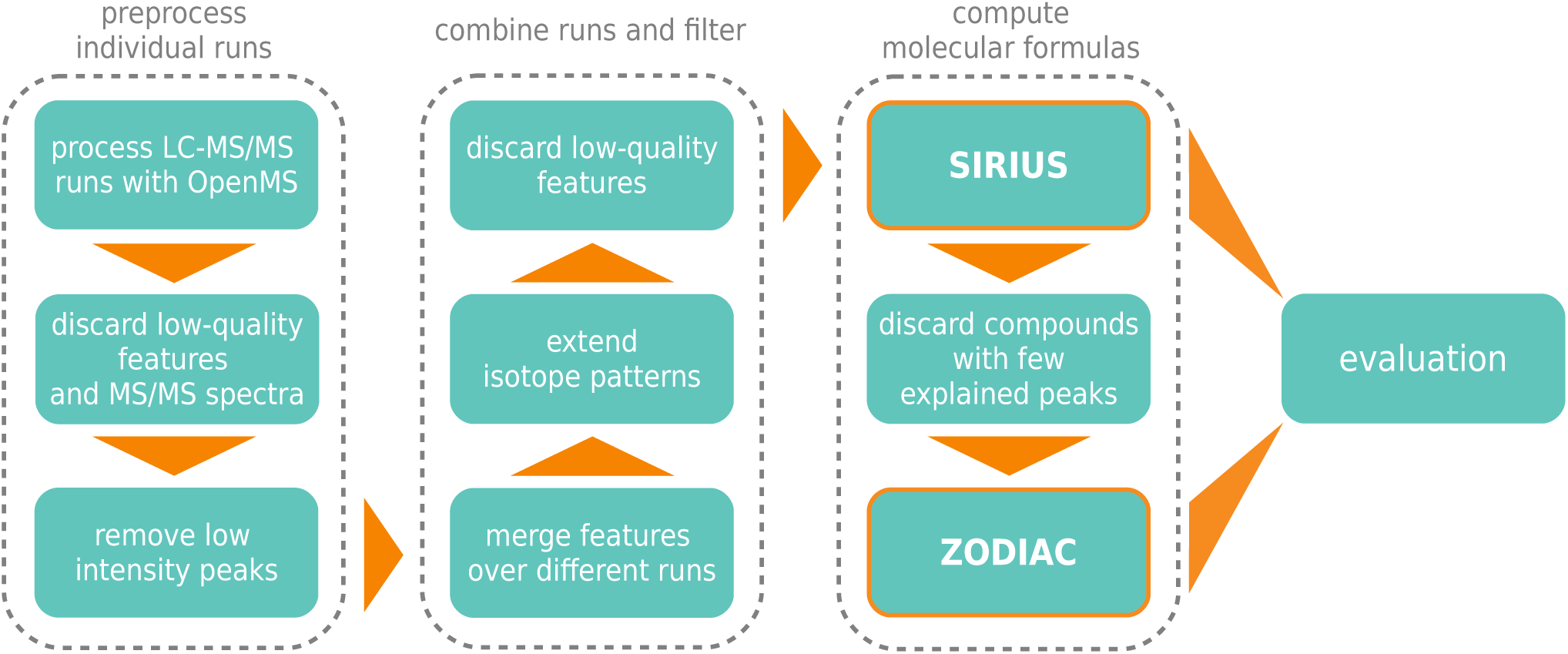
ZODIAC processing and evaluation workflow. 1) Each LC-MS/MS run is processed individually; input mzML/mzXML files are processed using OpenMS, performing feature and adduct detection and producing files in SIRIUS input format. Resulting features combine MS1, MS/MS and adduct information. 2,3) Filtering is performed on feature, MS/MS and peak level. 4) Similar features are merged between different runs using hierarchical clustering; MS/MS are combined and a best isotope pattern is selected per feature. 5) Missing isotope peaks are searched in MS1 spectra to extend isotope patterns. 6) A final feature filtering step is performed; the remaining features are considered as compounds. 7) SIRIUS is executed. 8) Compounds with few explained peaks are discarded, since a badly explained MS/MS spectrum indicates low quality. 9) ZODIAC is run on the remaining compounds. 10) SIRIUS and ZODIAC are evaluated on the same set of compounds.

**Supplementary Table 2:**
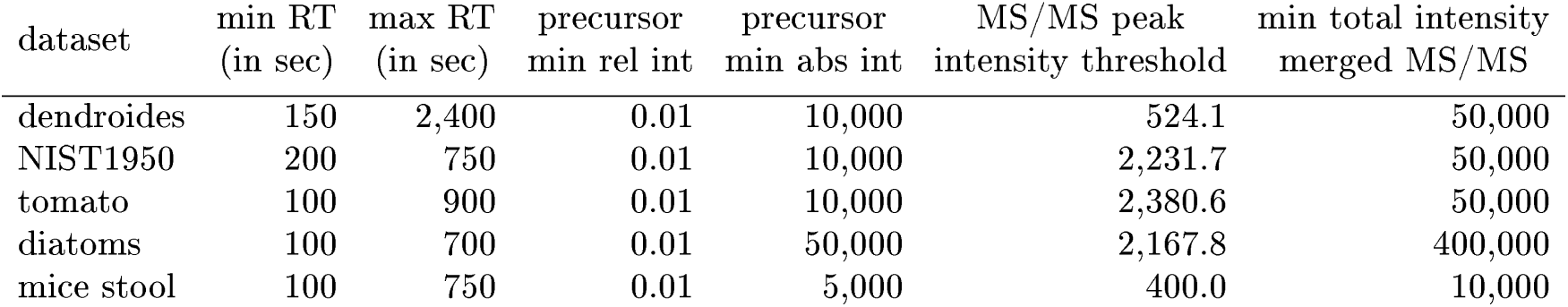
Parameters used to process and filter LC-MS/MS runs. Features were filtered by retention time (min RT, max RT) and minimum relative and absolute intensity of the precursor peak. MS/MS peaks below an intensity threshold were removed. MS/MS spectra were merged over different LC-MS/MS runs and discarded if total intensity was below an intensity threshold.

**Supplementary Figure 8:**
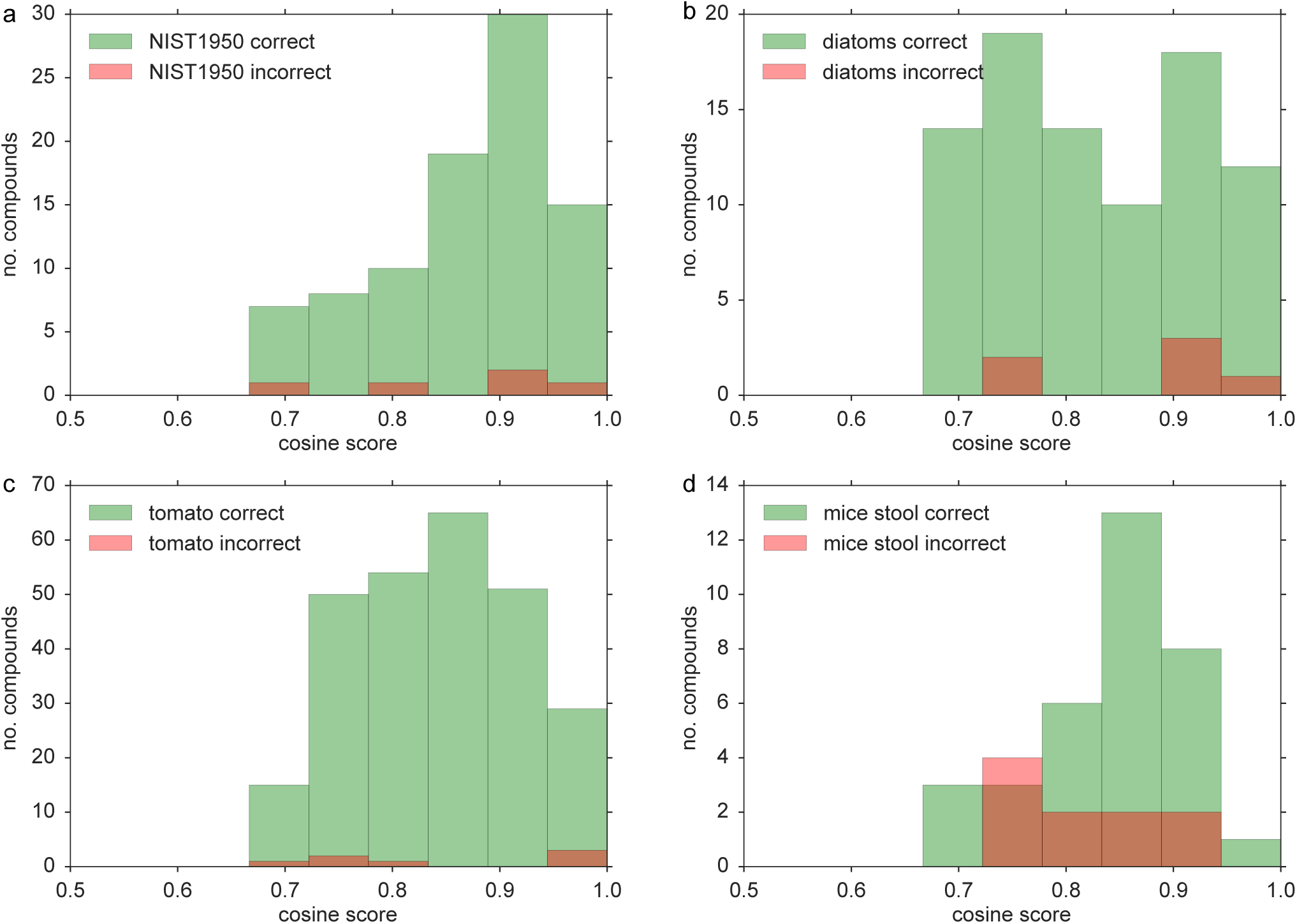
ZODIAC assignments vs. cosine scores of the ground truth. For four datasets, we can only evaluate ZODIAC against a “ground truth” established by spectral library searching. Potentially, some ground truth molecular formula are wrong, and ZODIAC might have found the correct molecular formula which we wrongly assign as incorrect. We expect that database hits with relatively low cosine score are incorrect more often. We have plotted the cosine score for correct and incorrect ZODIAC molecular formula assignments for NIST 1950 (a), diatoms (b), tomato (c), and mice stool (d). We *do not observe* a noteworthy difference in the two distributions; instead, correct and incorrect annotations appear to be distributed across all cosine scores. This does not mean that all library hits are correct, but that incorrect library hits are most likely to be found both for ZODIAC correct and incorrect assignments.

**Supplementary Figure 9:**
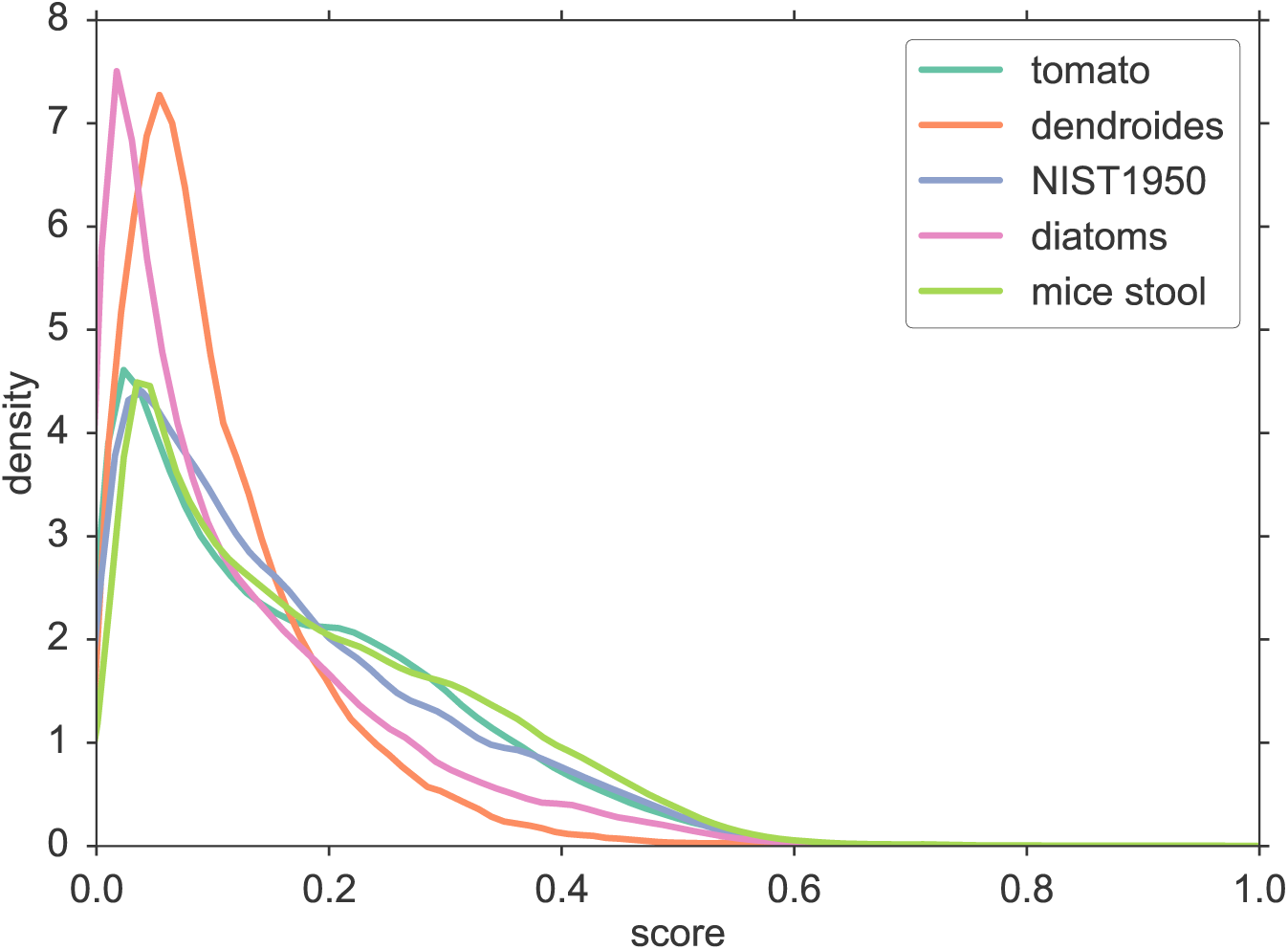
Distribution of fragmentation tree similarity scores. For each dataset, kernel densities were estimated using 100,000 sampled scores. Scores *s*(*u, v*) were computed as described in equation 6 in Section Materials & Methods.

**Supplementary Figure 10:**
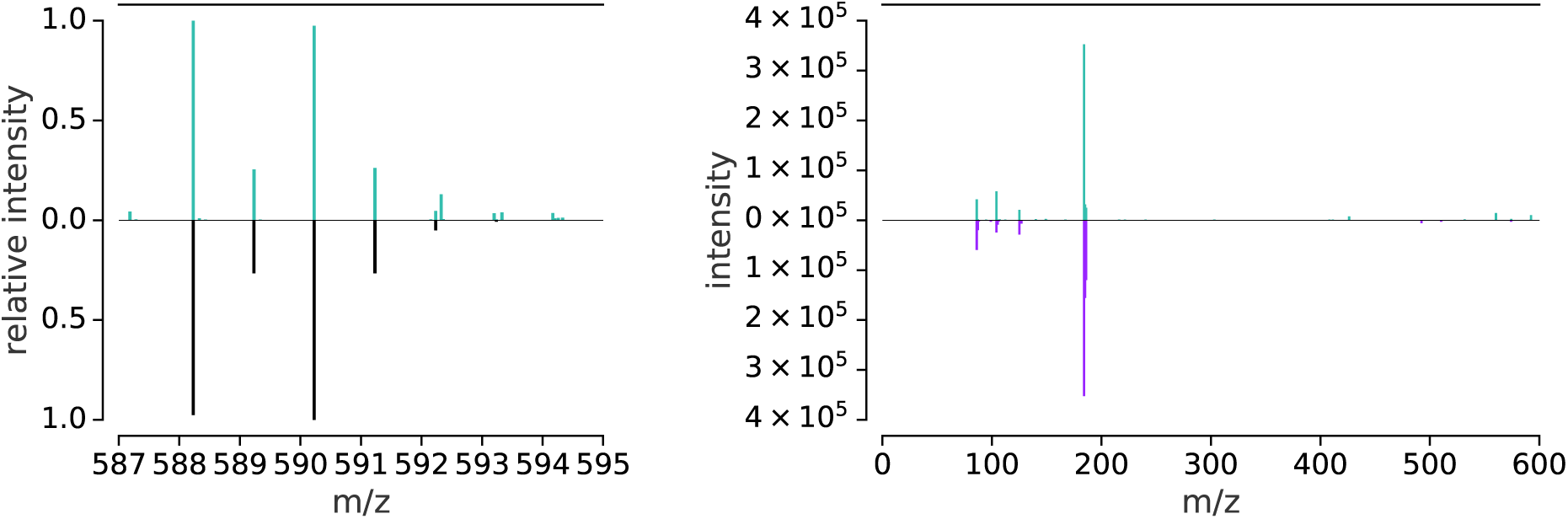
Spectra of a novel bromine-containing compound in the diatoms dataset. (left) Mirror plot of measured against simulated isotope pattern for the novel molecular formula C_24_H_47_BrNO_8_P in the diatoms dataset. The top part displays *m/z* 587 to 595 of the MS1 spectrum at retention time 505.18 sec. It was measured prior to the MS/MS spectrum targeting precursor *m/z* 592.325 and different from the MS1 in Fig. 3c, which the predecessor MS1 to the MS/MS spectrum targeting precursor *m/z* 588.230. The bottom part is the simulated isotope pattern for [C_24_H_47_BrNO_8_P + H]^+^. We see that close to the M+4 isotope peak, there is a more intense peak, presumably from a coeluting compound. Clearly, this coeluting compound can substantially affect the MS/MS spectrum. (right) Mirror plot of measured (top) against simulated (bottom) MS/MS spectrum for precursor M+4. Its intensity is one order of magnitude lower compared to the MS/MS spectrum of the M+2 peak and simulated intensities should be treated with caution.

**Supplementary Figure 11:**
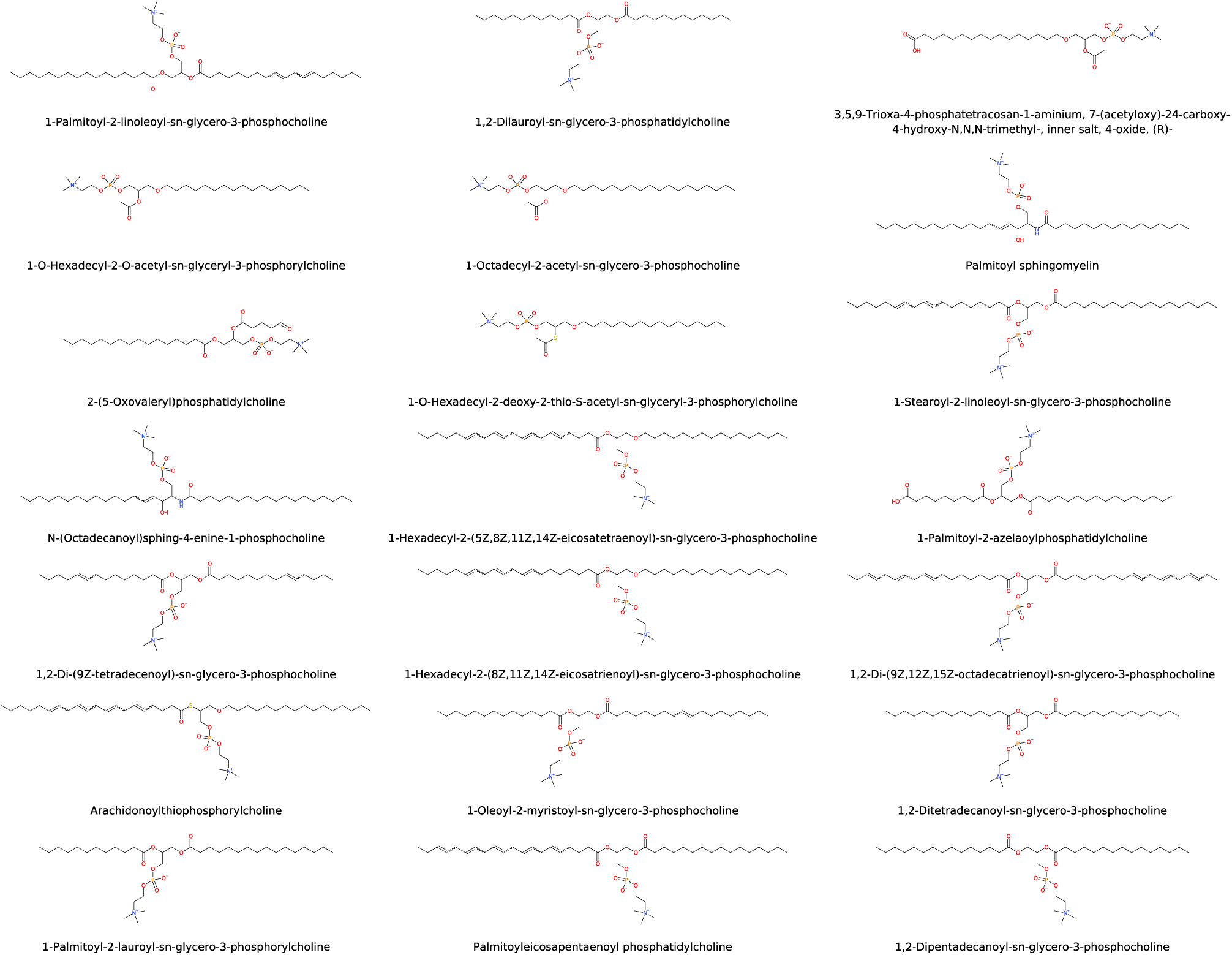
Structures of 21 NIST compounds matching to a novel compound in diatoms dataset. Structures are sorted left to right and top to bottom by cosine score to the query spectrum. All structures are phosphatidylcholines. Corresponding spectra are displayed in Supplementary Fig. 12.

**Supplementary Figure 12:**
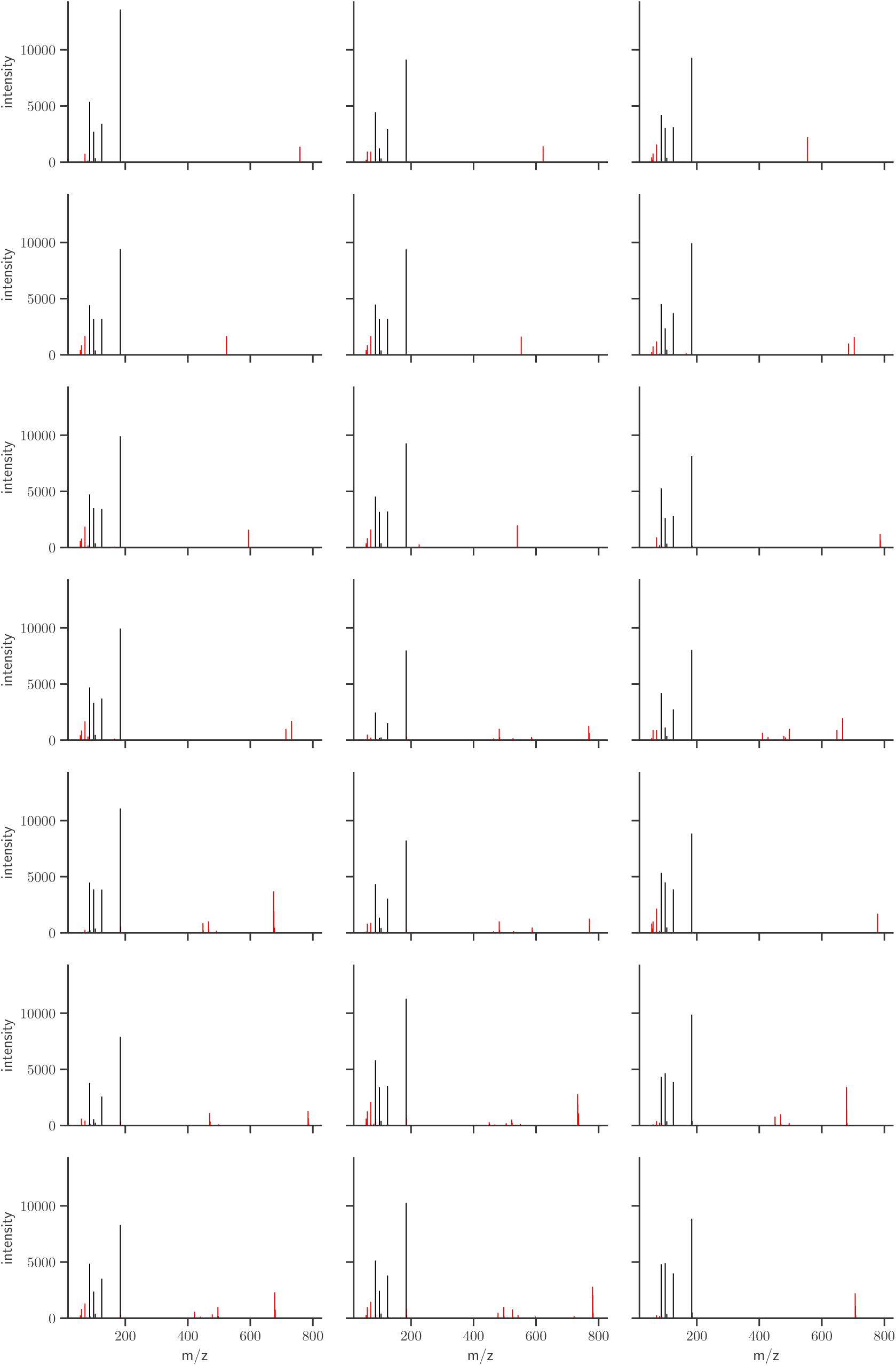
Spectra of 21 NIST compounds matching to a novel compound in diatoms dataset. Spectra are sorted left to right and top to bottom by cosine score to the query spectrum, with the lowest cosine score being 0.893. All spectra share a characteristic set of peaks; peaks matching to the query spectrum are displayed in black. The corresponding structures are displayed in Supplementary Fig. 11.

**Supplementary Table 3:** Novel molecular formulas. All molecular formulas are *absent* from the largest molecular structure databases PubChem^l7^ and ChemSpider^35^. Only molecular formula annotations with a minimum ZODIAC score of 0.999 are reported such that at least 95% of the MS/MS spectrum intensity is being explained by the SIRIUS fragmentation tree, and at least one molecular formula of the compound is connected to 20 or more compounds. There may be more than one hypothetical compound in an LC-MS run being annotated with one molecular formula, potentially corresponding to different isomers. For such cases, ‘# comp.’ is the number of hypothetical compounds being annotated with the given molecular formula, and ‘max score’ is the maximum ZODIAC score among these annotations. The corresponding compounds are given in Supplementary Table

| dataset | molecular formula | # comp. | max score | dataset | molecular formula | # comp. | max score |
| --- | --- | --- | --- | --- | --- | --- | --- |
| diatoms | C24H47BrNO8P | 6 | 1.0 | diatoms | C16H37N4O7 | 1 | 1.0 |
| diatoms | C22H47NO12 | 4 | 1.0 | diatoms | C16H39N4O6 | 1 | 1.0 |
| diatoms | C19H41NO11 | 3 | 1.0 | diatoms | C16H39N4O7 | 1 | 1.0 |
| diatoms | C21H45NO12 | 3 | 1.0 | diatoms | C16H39N4O8 | 1 | 1.0 |
| diatoms | C23H49NO13 | 3 | 1.0 | diatoms | C17H35NO11 | 1 | 1.0 |
| diatoms | C13H33N4O5 | 2 | 1.0 | diatoms | C17H41N4O9 | 1 | 1.0 |
| diatoms | C13H33N4O7 | 2 | 1.0 | diatoms | C18H39NO11 | 1 | 1.0 |
| diatoms | C17H41N4O7 | 2 | 1.0 | diatoms | C18H43N4O5 | 1 | 1.0 |
| diatoms | C17H41N4O8 | 2 | 1.0 | diatoms | C18H43N4O7 | 1 | 1.0 |
| diatoms | C18H43N4O8 | 2 | 1.0 | diatoms | C18H43N4O9 | 1 | 1.0 |
| diatoms | C23H49NO12 | 2 | 1.0 | diatoms | C19H39NO12 | 1 | 1.0 |
| diatoms | C24H49BrNO8P | 2 | 1.0 | diatoms | C19H45N4O10 | 1 | 1.0 |
| diatoms | C29H57N5O11 | 2 | 1.0 | diatoms | C19H45N4O7 | 1 | 1.0 |
| diatoms | C8H23N4O5 | 1 | 1.0 | diatoms | C19H45N4O8 | 1 | 1.0 |
| diatoms | C9H25N4O5 | 1 | 1.0 | diatoms | C19H45N4O9 | 1 | 1.0 |
| diatoms | C10H27N4O4 | 1 | 1.0 | diatoms | C21H49N4O10 | 1 | 1.0 |
| diatoms | C10H27N4O5 | 1 | 1.0 | diatoms | C21H49N4O9 | 1 | 1.0 |
| diatoms | C10H27N4O6 | 1 | 1.0 | diatoms | C22H51N4O11 | 1 | 1.0 |
| diatoms | C11H29N4O6 | 1 | 1.0 | diatoms | C23H53N4O11 | 1 | 1.0 |
| diatoms | C12H31N4O4 | 1 | 1.0 | diatoms | C24H49NO14 | 1 | 1.0 |
| diatoms | C12H31N4O6 | 1 | 1.0 | diatoms | C25H44NO9P | 1 | 1.0 |
| diatoms | C12H31N4O7 | 1 | 1.0 | diatoms | C27H57NO11 | 1 | 1.0 |
| diatoms | C13H33N4O6 | 1 | 1.0 | diatoms | C32H61N5O12 | 1 | 1.0 |
| diatoms | C14H33N4O6 | 1 | 1.0 | diatoms | C32H63N5O11 | 1 | 1.0 |
| diatoms | C14H35N4O6 | 1 | 1.0 | diatoms | C36H71N5O10 | 1 | 1.0 |
| diatoms | C14H35N4O7 | 1 | 1.0 | diatoms | C16H39N4O9 | 1 | 0.9995 |
| diatoms | C14H35N4O8 | 1 | 1.0 | diatoms | C18H43N4O10 | 1 | 0.9995 |
| diatoms | C15H35N4O5 | 1 | 1.0 | diatoms | C23H27O4P3 | 1 | 0.9995 |
| diatoms | C15H37N4O6 | 1 | 1.0 | tomato | C6H16N2O11P4 | 1 | 1.0 |
| diatoms | C15H37N4O7 | 1 | 1.0 | tomato | C27H60N10O4 | 1 | 1.0 |
| diatoms | C15H37N4O8 | 1 | 1.0 | tomato | C6H14N2O16P4 | 1 | 0.9995 |

Supplementary Table 4: **Compounds with a novel molecular formula**. Provided are the detailed information for compounds corresponding to the novel molecular formulas in Supplementary Table 3. All molecular formulas are *absent* from the largest molecular structure databases PubChem^l7^ and ChemSpider^35^.

THIS TABLE IS PROVIDED SEPARATELY IN FILE “compounds_with_novel_molecular_formulas.csv”.

Supplementary Table 5: **Manually annotated molecular formulas for compounds in the dendroides dataset**. These molecular formulas serve as ground truth for evalution of SIRIUS and ZODIAC.

THIS TABLE IS PROVIDED SEPARATELY IN FILE “groundtruth_dendroides.csv”.

Supplementary Table 6: **Spectral library hits for datatsets NIST1950, tomato, diatoms and mice stool**. The molecular formulas of these library hits serve as ground truth for evalution of SIRIUS and ZODIAC.

THIS TABLE IS PROVIDED SEPARATELY IN FILE “groundtruth_NIST1950_tomato_diatoms_micestool.csv”.

Supplementary Table 7: **List of input files used for evaluation of five datasets**. The included files in mzML/mzXML format correspond to LC-MS/MS runs which were used for evaluation. These runs are subsets of the data provided at MassIVE repository.

THIS TABLE IS PROVIDED SEPARATELY IN FILE “input_files.csv”.

